# Palaeoproteomics reveals bovine carcass processing with 6,000-year-old flint tools

**DOI:** 10.1101/2025.09.29.679186

**Authors:** Alexandra Burnett, Éva Halbrucker, Ian Engels, Rachel Siân Dennis, Isabelle De Groote, Dieter Deforce, Emanuela Cristiani, Philippe Crombé, Maarten Dhaenens

## Abstract

Lithic tools are the most abundant cultural artefacts found at prehistoric archaeological sites. Through their study, we can understand essential activities such as hunting, processing of animal carcasses and plant materials, and working of resources such as ochre and bone, as well as broader technological and cultural practices. For the first time, proteomics was applied alongside use-wear and optical microscopy residue analyses, enabling the identification of bovine, plant, and human proteins on faceted tools from the Mesolithic-Neolithic site of Bazel-Sluis, Belgium. This retrieval of identifiable protein from an area with poor organic preservation demonstrates wide-reaching potential for archaeological research, opening a new avenue of investigation into prehistoric lifeways and adding to the growing corpus of evidence accessible through palaeoproteomics. A workflow for future analyses is suggested, based on our integrated sampling strategy requiring no additional tool manipulations.

## Introduction

Lithic tools, used for a wide range of subsistence activities throughout prehistory, dominate material culture assemblages across Europe. Differences in the shape and size of tools, and the raw materials used for their manufacture, are broadly reflective of transitions in the actions performed with lithic tools, the materials on which the tools were used, and the communities producing and utilising lithic tools. While mostly studied in terms of typological, technological and geologic qualities, the application of microscopic and molecular analyses can extend our understanding of lithics and uncover traces of the materials with which the tools came into contact, adding also to our understanding of tool function and the activities performed at prehistoric sites.

Use-wear analysis is essentially the interpretation of damages to tool surfaces generated through tool use. The characteristics of use-wear traces such as edge damage, polish, and striation are indicative of the type of material with which the tool was in contact, and the motion of use. The interpretation of these traces is primarily dependent on experimental reference collections, however, some traces are less specific or interpretable than others, such as type 10 and 23 polishes^1^. As some materials produce more variable or less distinctive or developed traces than others, use-wear analysis inherently involves bias towards the identification of harder or more abrasive contact materials, contact materials which leave specific traces, and materials that were processed repeatedly. In most use-wear studies, soft material-related traces are therefore probably underrepresented.

Optical microscopy residue analysis is applied to microscopic remnants of materials which adhere to tool surfaces, and relies on unique visual and morphological characteristics to identify organic and inorganic substances^2^. Plant remnants are identifiable on the surfaces of ground stone tools as phytoliths and starch grains^3^, and some studies have also incorporated the broad identification of plant residues via simple staining tests^4^. However, there are relatively few studies in which plant remains have been identified on flaked tools^5^. These micro-residues are invaluable for determining tool use and processing activities. The identification of organic materials is sometimes hampered by poor preservation which alters the characteristics of the residues, while plant remains in particular may be suspected to arise from soil contaminations, culminating in less accurate identifications^2,6,7^. Inorganic residues, including some adhesive substances, preserve better than organic residues, and especially in the case of exotic or processed materials such as ochre, are more readily linked to tool use^7–11^.

The combination of use-wear and residue analysis techniques, comprising a holistic functional analysis, allows functional residues and environmental residues to be distinguished based on their relationship to wear traces, providing the most reliable biography of the tools^2,6–11^. This is essential because residues derived from hafting processes or contact with other substances in sediments may otherwise confuse interpretations of tool use. The development of increasingly high-resolution observational techniques has seen functional analysis grow into a more systematic and effective technique, encompassing optical microscopy, histological staining, and scanning electron microscopy (SEM)^12^.

Prompted by the development of new residue testing methods for forensic science^13^, a great number of studies aiming to detect traces of animal blood proteins on stone tools were conducted from the 1970s into the 2000s^1,14–29^. Such studies were commonly predicated on presumptive testing to identify reactions with the haem group of blood, followed by more targeted immunological, crystallographic, or electrophoretic analyses, often with some level of taxonomic identification. However, both presumptive and immunological tests demonstrated positive responses to chlorophyll, vegetable and bacterial peroxidases, modern animal faeces, and metal cations in soils^1,16,19^. Furthermore, both poly- and monoclonal antibodies are reactive to protein targets belonging to other species, including beyond the family level. One study damningly found that for cross-over immune-electrophoresis (CIEP), one of the most popular methods ^1,23,26,29–32^, commercial laboratories produced accurate species determinations in only 37% of cases^31^. Outcomes for CIEP-based plant analyses were no better^30^. These multi-step testing processes could also be highly time-consuming and required the use of a plethora of specialised reagents. Even excluding presumptive tests, Craig & Collins^33^ describe enzyme-linked immunosorbent assays (ELISA) protocols taking from 36 to over 102 hours, while Tuross, Barnes & Potts^22^ describe a protocol time of over 57 hours. To identify blood residues (absent species determinations) on 90,000-year-old tools from Tabun Cave in Israel, Loy and Hardy^19^ carried out a three-step analysis using Ames Hemastix colorimetric tests for myoglobin and haemoglobin, antibody dot-blot (gold immunoassay) testing with silver stain enhancement to identify immunoglobulin G (IgG), and finally high-pressure liquid chromatography (HPLC) to validate the presence of haemoglobin and serum albumin. In their CIEP analysis of Neolithic stone tools, Högberg and colleagues^1^ used 27 different antisera and obtained identifications with low taxonomic resolution – to the Clupeidae (herring) family and to percoids, and to the Cricetidae and/or Muridae families, arguably making the latter an assignment to the Muroidea superfamily.

Non-immunoassay techniques like haemoglobin crystallisation^14,17,25^ and sodium dodecyl sulphate-polyacrylamide gel electrophoresis^15^ (SDS-PAGE) were similarly flawed. Degraded proteins produce ‘smearing’ during electrophoresis which prevents identification by comparison to reference standards; plus some stains cross-react with soils^15^. Crystallography experiments demonstrated that degraded haemoglobin is incapable of forming crystals and that the findings of archaeological crystallisation experiments could not be replicated^18^. HPLC, another non-immunoassay technique, is the closest relative to modern liquid chromatography tandem mass spectrometry (LC-MS/MS) used for proteomics, but then required laborious manual spectra interpretation to identify few targets. Throughout the popularity of protein residue testing, many publications highlighted significant issues with its reliability^16,32,34–36^ (see ^37^ for a particularly thorough overview).

Following advances in mass spectrometry technology, protein analysis has developed into proteomics, wherein proteomes – the entire suite of proteins present in a sample, tissue, or organism – are analysed simultaneously, typically by LC-MS/MS. In this technique, peptides (digested proteins) are separated out along a gradient of fluid solvents, after which the masses of peptides are measured, and those meeting certain criteria for intensity, mass-to-charge ratio (*m/z*) and retention time, are blasted into smaller ‘daughter’ ions whose mass is also measured. The sequence of the peptides is then reconstructed based on the known masses of their 20 constituent amino acids (AAs), and allows them to be matched to the sequences of proteins in a database. This method of ‘shotgun’ proteomics allows every peptide to be acquired without prejudice, so that plant, animal, and microbial proteins can be analysed within a single sample. As only the primary structure is measured, and because non-tryptic breakages and post-translational modifications induced by taphonomy can be accounted for, LC-MS/MS can readily be applied to degraded samples. This powerful technique has been used to identify legumes, grasses, and dairy or meat/blood proteins from goat, sheep, bovids and cervids in Neolithic Anatolian pottery^38^, as well as for the characterisation of oral bacterial and human proteins from medieval Danish dental calculus^39^. These studies demonstrate that not only can palaeoproteomics – proteomic analysis of historical, archaeological, palaeontological, or cultural heritage samples^40^ – enable us to identify species, but also tissues, and states of disease and health. Furthermore, peptide sequence-based data reliably enables the construction of phylogenetic relationships which concur with or extend known taxonomic relationships^41^. As well as the utility of LC-MS/MS for taxonomic identification and tissue differentiation in simple and complex samples, proteomics also boasts greater sensitivity than immunoassay or chemical methods: ELISA is stated to detect less than 100 nanograms of protein per antibody type, and CIEP 10000 nanograms of a single protein^42,43^; meanwhile the MS/MS instrument used in this study is reportedly capable of quantitative analysis of 4600 proteins (43000 peptides) from 200 nanograms of a K562 cell lysate^44^. While LC-MS/MS has successfully been applied to a range of archaeological artefacts and organic remains, including paintings^45^, pottery^38^, bone^46,47^, enamel^48^, and mummified tissues^49^, its potential to resolve past issues of lithic protein analyses and investigate proteins bound to archaeological stone tools has yet to be realised.

The lithic artefacts analysed in the present study originate from a prehistoric wetland site, situated along the western bank of the Scheldt river in NW Belgium, close to the city of Antwerp (51°08’09”N, 4°19’23”E). Bazel-Sluis, excavated in 2011^50^, yielded artefact scatters consistent with a prehistoric settlement, including lithic artefacts^51^, pottery fragments^52,53^, faunal remains, charred hazelnut shells and cereal grains^54^, and charcoal^50,55^. These finds were recovered from a fairly homogeneous layer of fine sand, which was covered by organic clay and peat from ca. 3625-3390 cal BC onwards^56^. Peri-marine alluvial deposits with a clay-based composition subsequently covered the peat.

Two major occupation phases, separated by a period of non-habitation, have been described on the basis of thorough radiocarbon dating^58^, as well as ceramic^52,53^ and lithic^57^ artefacts. The first of the two occupations relates to an Early-to-Middle Mesolithic settlement in the 8^th^ millennium cal BC; in the second major phase, a number of Mesolithic and Neolithic cultural groups are represented from ca. 5200-3500 cal BC. These groups include Late Mesolithic hunter-gatherers, the Final Mesolithic-to-Early Neolithic Swifterbant culture, and the Middle Neolithic Michelsberg culture. However, it is difficult to ascribe individual settlement remains to either major phase, as a low sedimentation rate caused the finds to become somewhat intermingled ^59^. The intermixing of lithic artefacts could be “resolved” to a certain degree thanks to the preservation of a latent stratigraphy. Detailed analysis of the vertical distribution demonstrated a gradual increase in age of the artefacts with depth, the deepest artefacts belonging to the first occupation phase. This vertical “sorting” of artefacts could be linked to continuous and intense bioturbation of the soil, most likely caused by small burrowing animals such as moles and earthworms^59^.

The present study focuses on a specific tool-type from the second occupation phase, known as the faceted tool. Faceted tools consist of large flakes, debris fragments, discarded cores, and blocky pieces that cannot be clearly identified. Of these, denticulates, faceted flakes and debris are only partially shaped, while polyhedrons are fully shaped, by the crude removal of <20mm flakes. Faceted tools often feature wear traces visible to the naked eye, such as pecking and crushing along their ridges, and impact points. Unknown before ca. 5300 cal BC, this tool-type was introduced by the first farmer-herders belonging to the *Linearbandkeramik* Culture (LBK) who settled in the loess area approximately 80 km to the south of Bazel^60^. The use of faceted tools increased considerably during the course of the 5th millennium cal BC among subsequent Neolithic cultures, such as the Blicquy/Villeneuve-Saint-Germain culture^61^. By the mid-5^th^ millennium cal BC, faceted tools also appeared within indigenous hunter-gatherer contexts of the Swifterbant Culture within the Scheldt basin^51,62^. This is considered proof of increased contact and exchange between both communities, which is also supported by other evidence, such as the appearance of indigenous pottery^52,53^, cereal grains^54^ and domesticated animals^55^. As such, faceted tools constitute an important element in the study of Neolithisation along the Atlantic coast of NW Europe, situated just beyond the agro-pastoral frontier of the first farmer-herders.

In this study, we apply optical microscopic inspection of residues, use-wear, and proteomic analyses to 31 faceted lithic tools from Bazel-Sluis, which were previously subject to a typo-technological and raw material study^62^. Radiocarbon analysis of visible residue adhering to a tool surface provides a date for the use of the lithics, as well as demonstrating the antiquity of the preserved residues. The tools underwent an integrated analytical procedure, whereby the addition of proteomics to the workflow utilised waste cleaning fluid, non-destructive swabbing, and re-use of other samples. The results are used to reconstruct the motions of tool use, identify materials with which the tools came into contact, define the characteristics of the adherent residues, and to determine the taxa on which the tools were used.

## Results

### Use-wear traces

Thirty lithic tools were analysed for this study, on which 33 used zones (UZ) were interpreted [table **1**, **S2**]. The most abundant use-wear traces are related to animal processing (UZ=26). Of these, most traces (UZ=12) are related to contact with hard matter (e.g. bone, cartilage, antler) in a dynamic motion, requiring intensive movement [figure **1b**, **S1b**]. It was not possible to identify the exact action or specific contact material responsible for these traces through comparison with experimental tools. Based on partial overlap with traces from experimental tools used on fresh and slightly burnt bones^62^, the tools were likely used in a crushing, pulverising action in contact with fatty bone, probably also meat, tendons, skin and marrow [table **S1**, **S2**]. These use wear traces exhibit developed to well-developed characteristics. They are marked by smooth, flat, greasy, and bright polish localised primarily along the edges, occasionally extending into the background with mixed directionality. These polish patterns are consistently associated with extensive edge scarring across multiple layers, with pronounced battering evidenced by concentrated chipping along edges and ridges, and heavily rounded edges.

**Figure 1:**
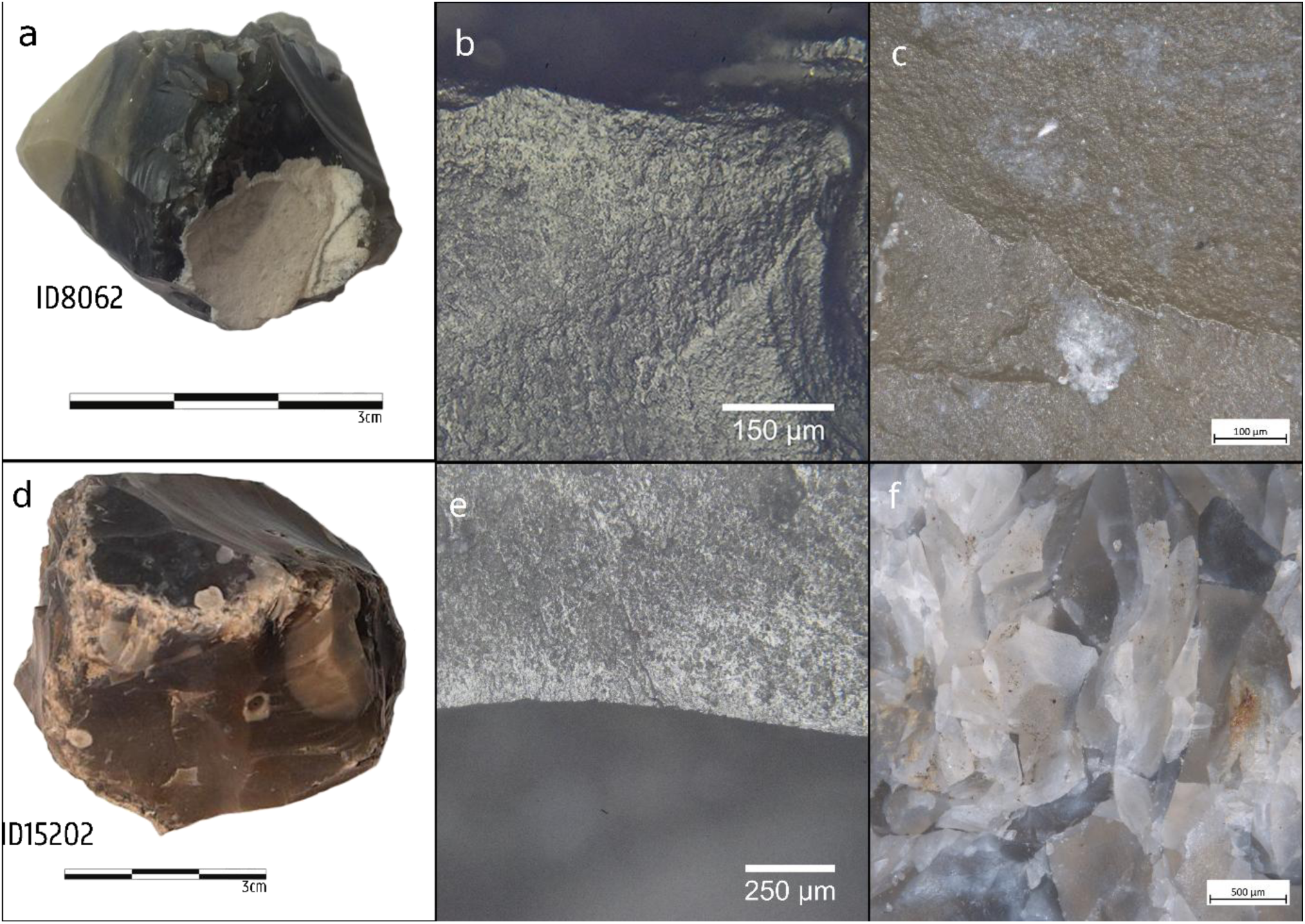
Bazel-Sluis faceted tools photographed with use-wear traces and visible residues: a) Tool 8062 b) traces of hard animal matter processing in a dynamic motion with rough and greasy, bright, flat and pitted polish with mixed directionality, rounded edges c) white, fatty, amorphous matter spread in the background d) Tool 15202 e) traces of medium hard animal matter processing in a dynamic action with rough and greasy, flat, bright polish in a transversal directionality f) beige-brown fatty, amorphous matter in use scars.

**Table 1:** Contact material and motion of tool use identified at used zones on faceted tools.

| **Contact Material** | **Motion of use** |  |  |  |  | **Total** |
| --- | --- | --- | --- | --- | --- | --- |
|  | Non-specific | Dynamic | Longitudinal | Transversal | Diagonal |  |
| **Animal** | **1** | **17** | **1** | **6** |  | **25** |
| Hard tissue |  | 12 |  |  |  | 12 |
| Medium tissue |  | 4 |  | 1 |  | 5 |
| Soft tissue |  | 1 |  |  |  | 1 |
| Hide dry |  |  |  | 1 |  | 1 |
| Hide fresh |  |  |  | 2 |  | 2 |
| Hide indeterminate |  |  |  | 2 |  | 2 |
| Meat/bone |  |  | 1 |  |  | 1 |
| Indeterminate | 1 |  |  |  |  | 1 |
| **Non-specific** | **1** |  |  |  |  | **1** |
| Abrasive | 1 |  |  |  |  | 1 |
| **Inorganic** |  |  |  |  | **1** | **1** |
| Hard stone |  |  |  |  | 1 | 1 |
| **Indeterminate** | **4** |  |  |  |  | **4** |
| **Total** | **6** | **17** | **1** | **6** | **1** | **31** |

Four used zones (Tools 8013, 9232, 15202, 16261) were interpreted as resulting from dynamic motion on medium-hard animal material [figure **1e**]. The traces are very similar to those on hard animal material but are more greasy, less smooth, and somewhat less developed. We interpret that these tools were used in a very similar way, on smaller game or less bony anatomy. One used zone had traces consistent with scraping motion on medium-hard animal material. Traces include rounded edges, rough to smoothening, greasy, flat, and bright polish banded along the edge on both dorsal and ventral sides with transversal directionality.

Five tools (7976, 10935, 13887, 18423, 18556) show traces related to hide scraping: two on fresh hide, one on dry hide, and two from hide in an indeterminate state. Traces include rounded to very rounded edges, with minimal scarring, while the polish is flat, and bright. In case of the fresh and indeterminate hide, the texture of the polish is rough and greasy, and in case of dry hide, it is rough and matt. On one of the tools for which the hide state is unknown, resharpening of the used edge is present. All tool surfaces presented collagenous or white indeterminate visible residues.

One tool (13552) was interpreted as having been used on soft animal matter in a dynamic motion on a hard surface. The polish is greasier and rougher than that generated by medium and hard animal material, but the edge scars are invasive and overlapping.

One used zone (tool 7634) features marks characteristic of contact with both meat and bone in a longitudinal cutting motion. The same tool also exhibits traces of dynamic action on hard animal matter [table **S1**]. The used zones of the two different types of traces are separate; the butchering traces are further from the edge and the edge scars. The polish is very greasy and rougher than that observed for bone alone.

One tool (17679) shows weakly developed indeterminate animal-related traces. They include overlapping invasive scalar edge scars on slightly rounded edges, and smoothening, greasy, flat and bright polish. Overlapping invasive edge scars indicate an intensive dynamic motion.

One tool (5258) exhibits well-developed nonspecific abrasive traces on a limited area of the tool with very smooth, matt, flat, bright polish with a lot of striations with random directionality. The same tool exhibits weakly developed unspecific traces connected to large scars and pulverising traces on several other areas.

Only one tool (11208) presents traces related to inorganic material, more specifically hard stone. Overlapping edge removals cover most of the surface of the tool, which has rounded to very rounded edges, and rough, matt, very bright, flat polish with diagonal directionality. The tool is thought to be a hammer stone for flint knapping or rejuvenating a (grinding) stone surface.

There are four used zones (tools 4694, 5258, 10068, 17665) for which the contact material could not be determined.

### Optical microscopy residue analysis

Microscopic inspection of Bazel-Sluis tool surfaces revealed remarkably consistent residue characteristics. Residues were detected on 24 of the 31 analysed tools [table **S1**]. Residue localisation and distribution were consistently associated with use-wear traces, and their specific characteristics were described. This suggests consistent patterns of tool use and a systematic association between wear traces and residues across the assemblage. The dominant pattern is white, fatty, amorphous matter found primarily in scars on the lithic tools. In general, the most common residue is white to beige or brown coloured, with amorphous fatty texture, though dry-looking residue also appears twice. The residue is typically accumulated and preserved in the (edge) scars, but in five cases also smeared in the background of the wear traces following the directionality of the executed movement [figure **1c**]. On two tools, feather barbules attributed to the Anatidae family were identified based on the appearance of their nodes and through comparison with reference material [figure **S1c**]. These barbules were connected to brown-white tissue and were smashed onto the surface of the tool, connecting with wear traces of dynamic use. Residues were also present on four tools for which the contact material was indeterminate based on the use-wear analysis. On one tool, unspecific white matter was described. These residues are best described as white or dark brown, fatty, amorphous matter mostly on the ridges and contact points, or in a band right behind the contact edge, or smeared on the surface and in edge scars. The brownish appearance of some visible collagenous fatty residues supports the interpretation of use-wear traces as resulting from rapid movement, since experimental work shows that very rapid and forceful motion can cause discolouration of the matter through friction between the tool and hard animal matter, especially bone [figure **1f**, see Supplementary Information for further comments]. Oxidation of iron-rich organic matter such as blood and bone marrow may also result in brown or reddish discolouration when exposed to air or prolonged friction.

Histological staining was performed on samples extracted from nine tool surfaces after macrowear and *in situ* analysis and before cleaning of the tools for microwear inspection (tools 4694, 7961, 8001, 10068, 11686, 15202, 16970, 17665, 18423). The binding of picrosirius red (PSR) dye, which appears as different shades of bright orange or green when applied to different types of collagen62, confirmed the presence of collagenous matter on all nine [figure **S1f**]. PSR-stained skin was identified on two tools (4694, 17665) and collagen on one more (tool 10068), on the basis of green and reddish-brown colour (respectively) and birefringence under cross-polarised light through transmitted light optical microscopy. A second round of staining analysis was carried out on sixteen more tools after they had been cleaned for use-wear analysis. Collagenous matter was stained from three of these (tools 8062, 16970, 18407), but it is plausible that some such matter was lost during the first cleaning process.

### Proteomic analysis

Samples were taken for proteomic analysis at three different stages of tool manipulation for use-wear and optical microscopy analysis of residues, hereafter referred to as ‘swab’, ‘soak’, and (microscope) ‘slide’ sample types. Peptides were extracted and searched against three reference databases: the entirety of SwissProt; the CollagenDB^64^; and blood, muscle, milk, and major seed proteins (the ‘CerealKiller’ search); all alongside common contaminants [see Methods]. A total of 380 proteins (2967 peptide sequences, 3534 peptidoforms; 595,308 PSMs) were identified across 60 samples from 19 lithic tools in three database searches [figure **S3**]. The ‘soak’ samples of tool sonication solution (*n*=44) yielded the greatest number of unique proteins and peptides, followed by the ‘swab’ samples taken from uncleaned tool surfaces (*n*=8), while the ‘slide’ samples representing a subsample of the sonication solution (*n*=8) produced the fewest protein and peptide identifications [figure **S2**]. The protein and peptide yield per tool for each sample type was highly variable, and was likely impacted by non-uniform sample quantities. Due to issues of protein similarity, each peptide-spectrum match (PSM) yielded identifications for up to 5 different taxa when combining search outputs, though most matched to proteins belonging to only one (74.5%) or two (16.2%) species. *Homo sapiens* (human) was the most dominant taxon in terms of unique protein identifications (*n*=171) across the dataset, though many of these are known skin contaminants, with keratins alone accounting for 42 protein identifications. *Bos taurus* (cattle) had the second-most unique proteins (*n*=40). *Mus musculus* (mouse; *n*=18) and *Ovis aries* (sheep; *n*=8) proteins were almost exclusively keratins, with high sequence similarity resulting in duplicate taxon hits.

Interestingly, numerous human proteins pointed to the presence of saliva on tool surfaces, including salivary biomarkers alpha-amylase 1A, cystatin-SN, and statherin^65^ [figure **S4, S5**]. Overall 92 proteins were found to overlap with those found in a separate analysis of modern saliva [figure **S4**, **S5**]. There were also a surprising number of plant peptides, from species such as *Anacardium occidentale* (cashew), *Prunus dulcis* (almond), and *Juglans regia* (walnut), which are not evidenced in Neolithic North-Western Europe [table **S3**]. *Zea mays* (maize) peptides are considered unremarkable and likely due to the known presence of corn starch on weighing papers used for residue transfer. Other plant species such as *Corylus avellana* (European hazel) and *Arabidopsis thaliana* (mouse-ear cress) are more ambiguous and may derive from contact with the tool surface during the Mesolithic/Neolithic or after excavation^54,66,67^. Many of the archaeologically plausible species fall into the Poaceae (grasses) family, which were the most well-represented in the proteomic data [figure **2**, table **2**, **3**]. A BLASTp search of all peptides for *Avena sativa* (oat; 24 peptides), *Corylus avellana* (*n*=21)*, Triticum aestivum* (wheat; *n*=6), and *Arabidopsis thaliana* (*n*=2) [**Supplementary Datafile**] revealed the presence of specific peptides for oat and hazel proteins (*n*=9, 3), as well as substantial overlap with peptides of *Alopecurus aequalis* (orange foxtail) and *Carpinus fangiana* (monkeytail hornbeam, a member of the birch family) respectively. No specific peptides were found for *T. aestivum* or *A. thaliana*, though one peptide was specific to the *Triticum* genus. However, due to limited evidence in individual tools [table **2****, 3**], and because database search strategies are more performant for the identification of proteins than for species, these taxa should be treated with a degree of caution. It is most likely that there are inaccurate plant taxonomic assignments resulting from incomplete protein databases – given this, it is difficult to estimate whether these are ancient or modern in origin [see also Supplementary Extended Discussion; figure **S8**].

**Figure 2:**
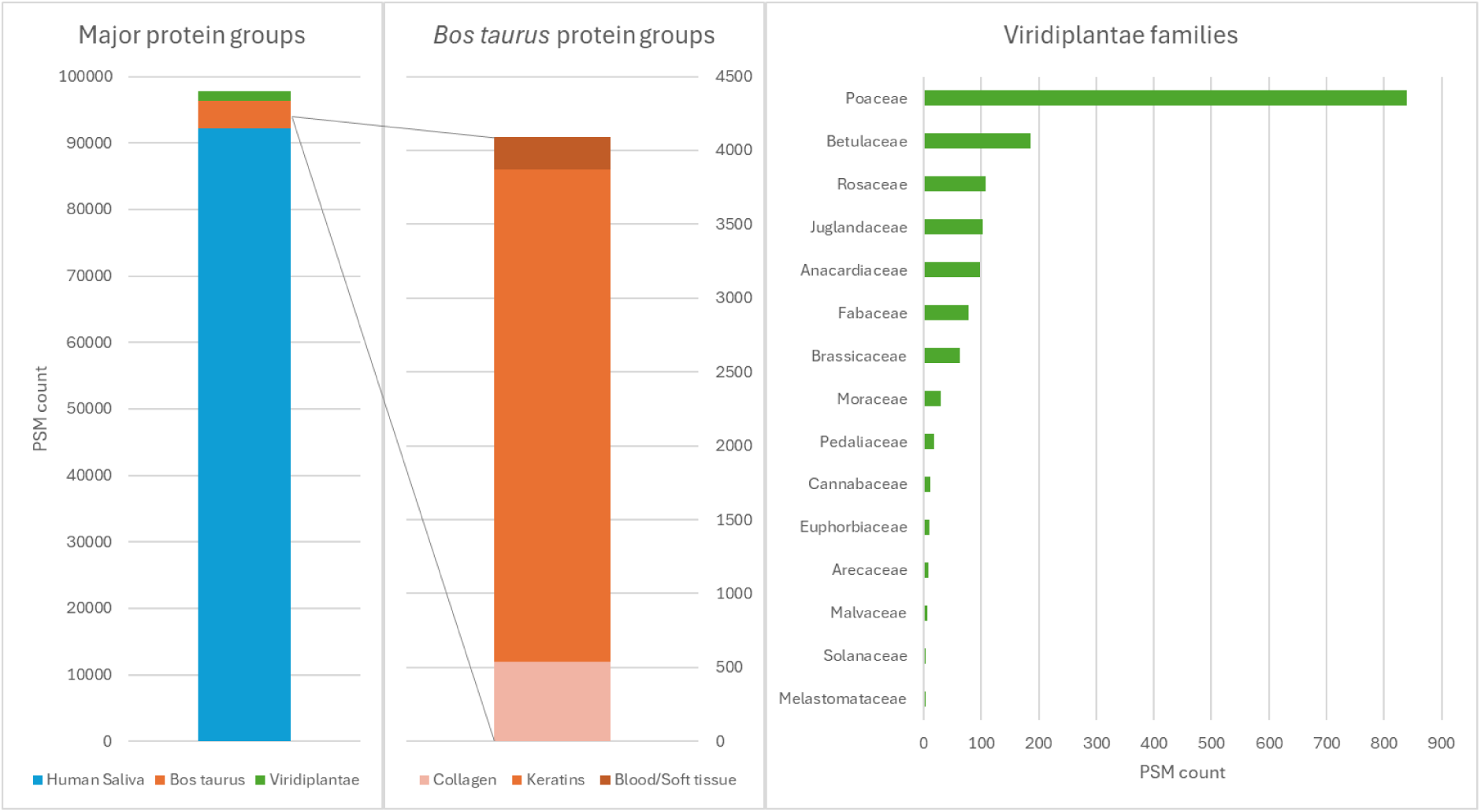
Overview of main protein groupings in the dataset. Left) human salivary protein PSMs dominate the proteomic profile of the lithic tools; Middle) bovine protein PSMs are largely attributable to keratins, followed by collagen type I and blood/soft tissue proteins; Right) Poaceae account for more than half of PSMs identified as Viridiplantae, including genera Avena, Triticum, Lolium, and Phragmites.

**Table 2:** Distinct peptide counts per species for tool 4694. To avoid duplication of query hits, counts for Homo sapiens and Bos taurus are derived from the SwissProt search, and all plant counts were taken from the CerealKiller search.

| Tool 4694 | Slide | Soak |  |  |  | Swab | Species Peptide Total |
| --- | --- | --- | --- | --- | --- | --- | --- |
|  |  | rep1 | rep2 | rep3 | rep4 |  |  |
| **Mammalia** |  |  |  |  |  |  |  |
| *Homo sapiens* | 202 | 233 | 181 | 171 | 209 | 270 | 416 |
| *Bos taurus* | 14 | 5 | 7 | 6 | 10 | 8 | 20 |
| **Viridiplantae** |  |  |  |  |  |  |  |
| *Arabidopsis thaliana* | 1 | 1 |  | 1 | 1 |  | 1 |
| *Avena sativa* | 3 | 1 |  | 3 | 3 |  | 3 |
| *Corylus avellana* |  |  |  | 1 |  |  | 1 |
| *Oryza sativa Japonica Group* | 1 | 1 |  | 1 | 1 |  | 1 |
| *Triticum aestivum* | 2 |  |  |  |  | 1 | 2 |

**Table 3:**
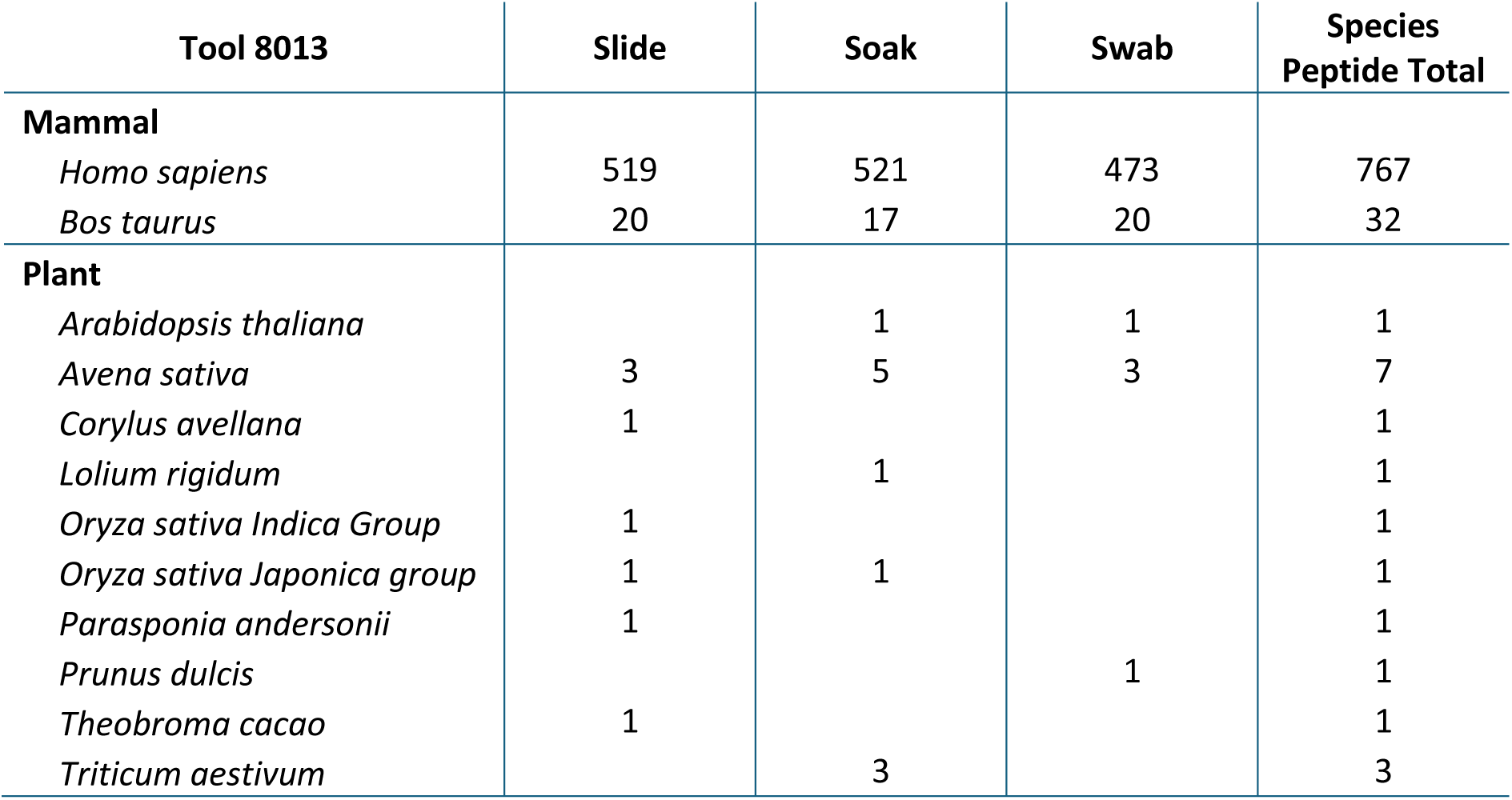
Distinct peptide counts per species for tool 8013. To avoid duplication of query hits, counts for Homo sapiens and Bos taurus are derived from the SwissProt search, and plant counts from the CerealKiller search.

| Tool 8013 | Slide | Soak | Swab | Species Peptide Total |
| --- | --- | --- | --- | --- |
| **Mammal** |  |  |  |  |
| *Homo sapiens* | 519 | 521 | 473 | 767 |
| *Bos taurus* | 20 | 17 | 20 | 32 |
| **Plant** |  |  |  |  |
| *Arabidopsis thaliana* |  | 1 | 1 | 1 |
| *Avena sativa* | 3 | 5 | 3 | 7 |
| *Corylus avellana* | 1 |  |  | 1 |
| *Lolium rigidum* |  | 1 |  | 1 |
| *Oryza sativa Indica Group* | 1 |  |  | 1 |
| *Oryza sativa Japonica group* | 1 | 1 |  | 1 |
| *Parasponia andersonii* | 1 |  |  | 1 |
| *Prunus dulcis* |  |  | 1 | 1 |
| *Theobroma cacao* | 1 |  |  | 1 |
| *Triticum aestivum* |  | 3 |  | 3 |

Mammalian collagens were identified from all but one tool, regardless of extraction type [table **S4**, **S5**]. Collagen type I constitutes over 90% of organic matrix content in bone^68^, and is comprised of one alpha-2 and two alpha-1 chains. Of eight species for which collagens were identified in the CollagenDB search, a number could be excluded on the basis of low number of peptides or taxonomic unlikeliness (*Apodemus sylvaticus, Chlorocebus sabaeus, Ictidomys tridecemlineatus, Myotis lucifugus, Phyllostomus discolor;* respectively wood mouse, green monkey, thirteen-lined ground squirrel, little brown bat, and pale spear-nosed bat); only *Bos taurus* collagen type I alpha-1 (COL1α1) and alpha-2 (COL1α2) chains, and *Sus scrofa* (pig/boar) COL1α2 showed consistent presence across all samples. A total of 39 peptide sequences belonging to *Bos taurus* collagens were identified, belonging to 49 peptidoforms. All *Sus scrofa* peptides (9 sequences, 11 peptidoforms) were identical to a subset of those assigned to *Bos taurus*.

To confirm this outcome, ClassiCOL^64^ analyses were performed. During this analysis, all peptidoforms from the database search were matched to all possible peptidoforms in the CollagenDB with identical molecular weights, to generate a more expansive list of potential peptide candidates (PPCs) for species classification. In this way, search engine annotation errors are accounted for by ensuring that the correct peptide annotations are present among the PPCs used by ClassiCOL to assign probable species scores. Four tools passed the minimum scoring and peptide number quality thresholds (>0.5 taxon score and >20 peptides for the top taxon). Tools 4694, 7961, and 8013 returned the *Bos* genus and *Bison bison bison* as equal top hits, while for tool 7961, *Muntiacus* (Muntjak deer) also shared the top score. Tool 5258 also passed the scoring threshold but returned both *Capra hircus* (goat) and *Odocoileus virgianus texanus* (Texas white-tailed deer) as top hits, with the *Muntiacus* genus close behind – a somewhat ambiguous ‘genetic mixture’ result [figure **S7**]. This may indicate the processing of either *Capra hircus* or a cervid absent from the database (such as *Capreolus capreolus* (roe deer)), or of both. *Cervus elaphus* (red deer) was discarded by the algorithm. Of those tools that did not meet the scoring quality thresholds, tools 8001, 13887, 17665, and 18423 also returned *Bos*/*Bison* or Bovinae as top hits. All other tools which failed to meet the quality threshold returned ambiguous results below the level of Pecora, while tools 16970, 18407, 18532, and 18556 had too few peptides to be classified at all.

Thirty-eight further *Bos taurus* proteins were identified in the dataset overall [table **S4**]. These include keratins type I cytoskeletal 28, type II cytoskeletal 71 and 73, type II cuticular Hb1 and Hb3, supported by 71 distinct peptides, 8 of which are unique to *Bos taurus* keratins among all proteins identified in the SwissProt search [figure **S6**]. There is otherwise significant overlap with human keratins (prevalent in the saliva extract), and a BLASTp search also demonstrated that *Ovis aries* is an equally likely source of the ‘unique’ keratin peptides, meaning that these could also be derived from clothing. Few *Bos taurus* proteins were present in all sample types. These include actin, cytoplasmic 1; alpha-S2 casein; elongation factor 1-alpha 1; kappa casein; and serine protease 1. Caseins are commonly considered to be dairy proteins but are also present in mammary tissue. Bovine alpha-amylase identifications were only given in the event that human amylase was absent in search databases. *Bos taurus* 14-3-3 protein zeta/delta and tubulin beta-5 chain were identified in all three database searches, but were not found in the microscope slide samples. Interestingly, serine protease 1 was identified more prevalently in the microscope slides than in swabs and soaked samples, across all searches, in a reversal of the overall trend of peptide and protein content for the different extractions [figure **S2**]. While this suite of cellular and plasma proteins may appear to indicate the presence of soft tissue residue on tools, the fact that a number of these constitute common contaminants means that these are not considered to be reliable evidence.

The number of human and bacterial peptides was highly variable between samples and extraction methods, with broadly more in the sonication water and swabs than the microscope slides – a trend which is likely reflective of the sampling strategy. Direct comparisons for tools 4694 and 8013 demonstrate no clear trends in plant, *Homo sapiens* and *Bos taurus* PSM counts between the three sample types [table **2****,3**]. High human counts are reflective of the presence of saliva and skin proteins resulting from extensive tool manipulations.

### Radiocarbon dating

Just one faceted tool yielded sufficient organic residue to perform radiocarbon dating, producing a date of 4473±261 BP (ETH-141995). Due to the large standard deviation, resulting from the very small sample size [see Methods], the date has a low precision, situating the artefact in the late 5^th^ to the very start of the 3^rd^ millennium cal BC (4226-2893 cal BC, 95.4% probability). Considering that the site was sealed by organic clay and peat from ca. 3625/3390 cal BC onwards and none of the 110 radiocarbon dates performed on animal bone or charred plant remains (cereal grains, hazelnut shells) is younger than 3500 cal BC^50,55,58^, the mostly likely age of the dated residue is late 5^th^ or first half of the 4^th^ millennium cal BC. This corresponds to the Michelsberg Culture occupation phase of the site, considered as the first fully agro-pastoral culture in the Scheldt basin.

### Integrated results

The outcomes of the optical microscopy analysis of use-wear and residue, and proteomic analyses concur well with one another, revealing that of the studied faceted tools, most were used to process animal carcasses during the Mesolithic-Neolithic occupation of Bazel-Sluis [e.g. figure **3**, **S8**]. For the first time, the application of shotgun proteomics has been used to identify the species to which animal residues on lithic tools belong, adding information to indications of collagen provided through visualisation and histological staining techniques. As it is not yet possible to distinguish between *Bos taurus* (domesticate cattle) and *Bos primigenius* (aurochs) proteomically, the identification of *Bos taurus* proteins should be considered as indicative of either species. While those tools whose results did not surpass the ClassiCOL quality criteria should be viewed as less reliable than those for which there were a greater number of supporting PPCs, it is notable that all four with a low-confidence single genus-level outcome were also attributed to bovids. In the case of Tool 5258 there is also potential evidence for goat proteins, but the equal scoring of goat and Texas white-tailed deer may also indicate the processing of roe deer, a close relative whose collagens are absent from available protein sequence databases [figure **S7**]. It is possible that re-searching of the data using experimentally constructed roe deer sequences would be useful to resolve this, but this is beyond the scope of the current study.

**Figure 3:**
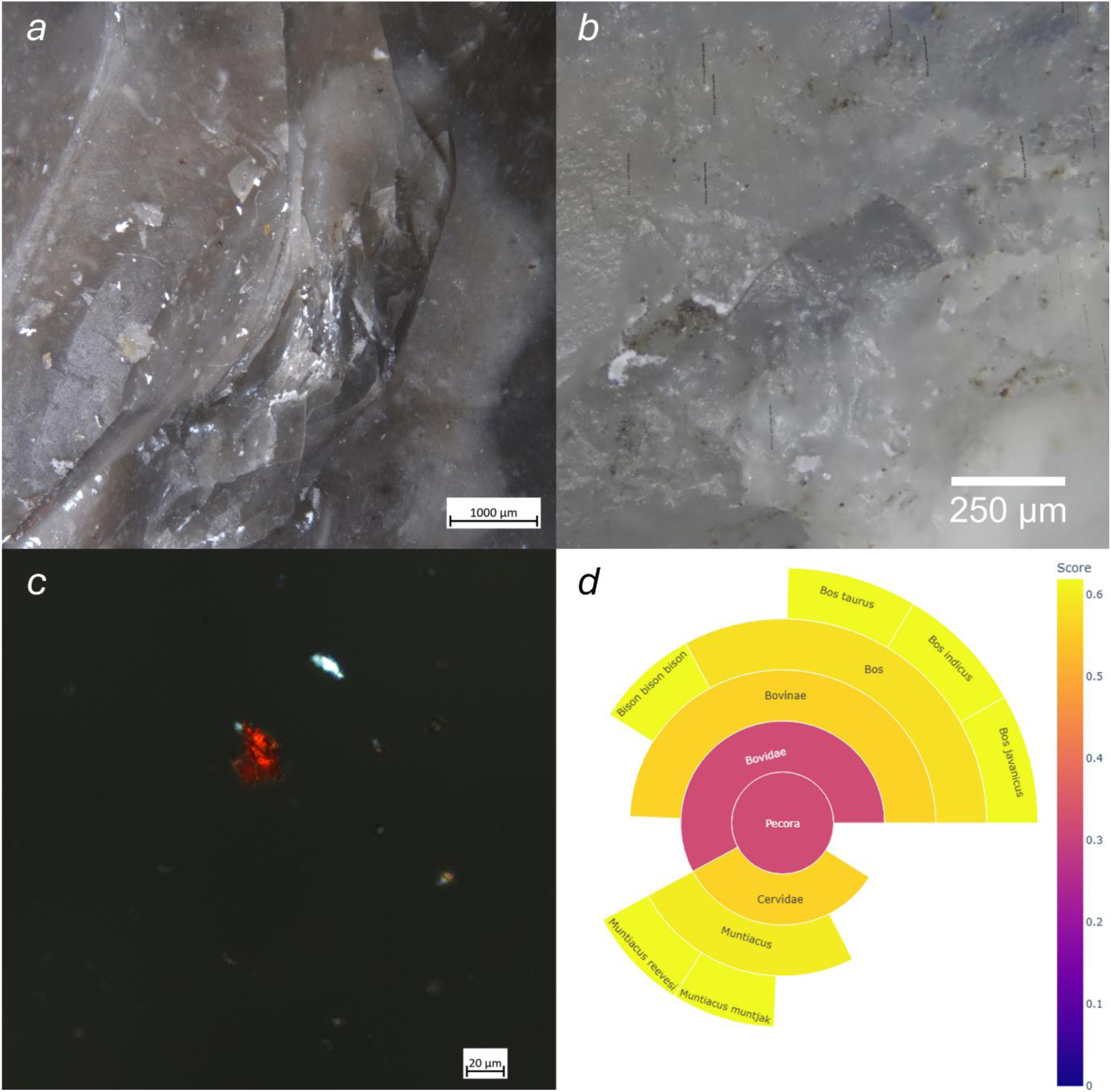
Multiple analyses of Bazel-Sluis lithic tool 7961 demonstrate consistent evidence for the processing of collagenous animal material: a) white, clumpy, dry matter concentrated in use scars, b) developed smooth, domed with flat parts, greasy, bright polish on the very rounded, but crushed edge of the tool with mixed directionality indicative of contact with hard animal material in a dynamic repetitive motion, c) collagenous tissue stained with Picrosirius red under cross-polarised light, d) extracted collagen residues are identified as Bovine in a ClassiCOL analysis.

Visible feather barbules were not represented in the proteomic results, though this is possibly due to the exclusion of barbules while sampling microscope plates. Additionally, the evidence for plant peptides across the assemblage is not well-mirrored by the microwear and visible residues, and in the case of tool 9232, traces of contact with harder animal tissues is incongruous with observed plant fibres [table **S1**]. It is as-yet unclear whether proteomics is able to detect otherwise inaccessible prehistoric plant residues, or whether these represent post-depositional contaminations.

Crucially, the result of the radiocarbon dating demonstrates the preservation of collagenous-appearing prehistoric residues on tool surfaces without significant contaminations which would otherwise produce an artificially young date.

## Discussion

This first use of LC-MS/MS for lithic tool-bound protein analysis represents a clear advance on prior methods of protein analysis, enabling identification of both plant and animal taxa through a non-labour-intensive analysis that furthermore does not require highly specialised reagents. In dealing with this particularly sparse proteomic data, we have endeavoured to avoid overinterpretation, carrying out confirmatory saliva and collagen analyses and applying conservative quality parameters. We have also carefully considered the occurrence of ‘analytically true, archaeologically false’ outcomes, which may arise due to sequence similarities and database deficiencies, or contaminations arising from the burial, excavation, and post-excavation environments. In doing so we have approached protein identifications as archaeologically false unless it is highly unlikely to be otherwise [figure **S9**].

The dominant presence of *Bos* proteins on the analysed faceted tools agrees well with the faunal spectrum of the site^50,55^. Among the 416 bone fragments that could be morphologically assigned to a species, 60% belong to aurochs/cattle. Wild boar/pig and sheep/goat represent resp. ca. 21% and ca. 5%. Among the exclusively wild game species, red deer predominates with ca. 9%, while other species (roe deer, beaver, hare, …) account for at maximum 1.5%. The use of faceted tools for crushing bone might be related to the extraction of highly nutritious marrow and/or grease, or the making of adhesives. When considering the amount of bone-crushing required for each activity, extraction of marrow/grease is the most labour-intensive, and as the bones of ruminants contain the highest proportion of marrow, bovids represent an ideal resource. Additionally, evidence for ruminant fats in Swifterbant and Michelsberg pottery from the Scheldt river basin, including Bazel-Sluis^69^, further evidences that ruminant carcasses were processed at Bazel-Sluis, as dietary components and possibly for the rendering of grease from crushed bones by boiling^70,71^. The apparent presence of human saliva might cautiously be attributed to *ad hoc* sample cleaning during excavation or tool examination. This may be especially true in moist, plastic sediments, which are not easily removed to determine shape, material and object type. This also provides a possible explanation for the presence of plant peptides which cannot plausibly be derived from a Mesolithic/Neolithic context, such as almond and walnut, that may represent remnants of food and drink consumed shortly prior to the recovery of the tools or during post-excavation handling. This hypothesis is of course highly speculative. The proteomic and microscopy evidence for plant residues should be viewed with some caution especially given the lack of observed plant-associated use-wear traces. It is possible that the plant residues represent later contaminations, however, the identification of plant macrofossils including *Triticum*, *Corylus avellana*, and *Prunus spinosa* (blackthorn, or sloe) at Bazel-Sluis^54^, fit well with the proteomic findings. Due to the time distance since excavation of the site, it was not possible to analyse soil samples from lithic findspots, which might otherwise have been useful to rule out contaminations from the burial environment. Another potential source of protein on tool surfaces is organic hafting materials, such as sinew^72^ and birch pitch^73^ softened through chewing. However, as we have no indication for hafting of faceted tools, this is an unlikely source of salivary or plant proteins. It is clear that further work is necessary to disentangle inaccurate database matches from potential plant contaminations so that richer information can be obtained in future analyses.

Use-wear evidence for inorganic substance processing, unique identifications through optical microscopy, and proteomic taxonomic insights underscore the necessity of multi-method approaches to capture the full breadth of information offered by archaeological lithics. It remains crucial that microwear evidence is used to link residue deposits with the use of the tools. The retrieval of proteinaceous residues from an area of the site with poor organic preservation is promising for the applicability of this method to sites with middling or poor preservation conditions across north-western Europe and beyond [Supplementary **Extended Discussion**]. The sampling strategies employed were all successful in retrieving informative peptides, evidencing that proteomic analysis can be applied to assemblages without necessitating additional sample manipulations. The current methodology can be applied to any tool that undergoes a cleaning process; the success of the microscope slide extractions is particularly promising for the analysis of microliths, whose small use edges would require little cleaning solution.

It is possible that a two-step cleaning process, in which a first sonication is used to remove superficial modern proteins and microscopic residues, and the second sonication is used to retrieve more strongly bound residues and proteins deposited deep in tool microcracks and depressions ^28^, might enable enhanced retrieval of ancient peptides (for example by diminishing ion suppression) [**figure 4**]. Concentration of wash solutions before protein extraction is likely to increase protein yield. Additionally, as sonication of tools in plastic bags may contribute to ion suppression by PEG contamination, testing of alternative cleaning vessels is desirable in future research.

**Figure 4:**
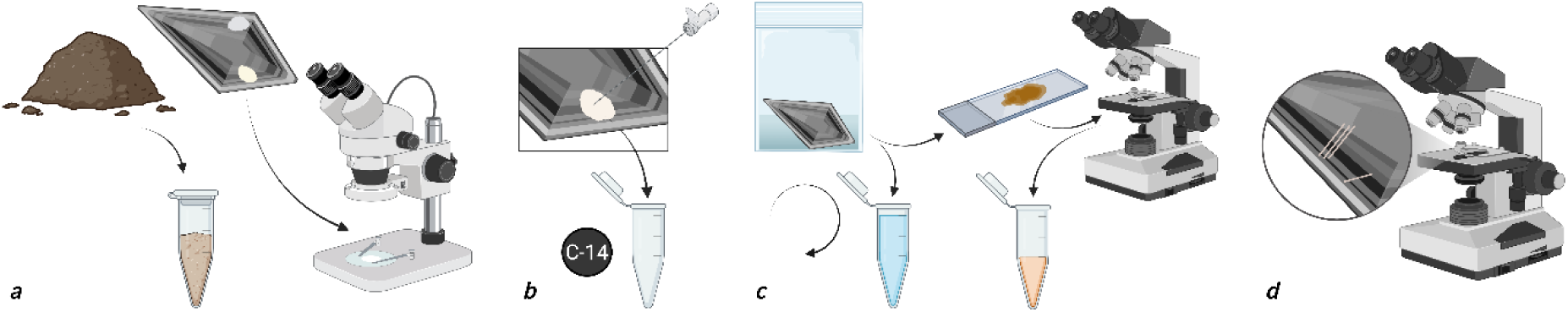
A suggested workflow integrating sampling of lithic residues for radiocarbon, proteomic, residue and use-wear analyses: a) soil from the lithic findspot is sampled as a proteomic control, and in-situ residues are directly recorded; b) large visible residues are sampled for radiocarbon dating; c) tools are sonicated, and the sonication solution collected for proteomics with a subsample used to prepare a slide for histological picrosirius red staining. After microscopic analysis the stained material is removed for proteomic analysis. Sonication is repeated in a denaturing solution to retrieve additional material for proteomics; d) the cleaned tool undergoes use-wear analysis.

In summary, this first proteomic analysis of lithic tools demonstrates congruence with the archaeological context and orthogonal analyses, revealing predominantly bovine processing and potential early domesticate plants at Mesolithic/Neolithic Bazel-Sluis. The relationship between the use-wear traces and the orientation, appearance, distribution, and localisation of residues, along with the radiocarbon dating, have been integral to marry the surface residue findings to the actual use of the tool, and will continue to be indispensable in future research. The success of this analysis at a findspot with conditions that are not conducive to organic preservation shows great promise for the application of proteomics to other lithic assemblages, while proposed methodological refinements should enable richer outcomes.

## Methods

### Use-wear traces

Thirty-one faceted lithic tools from the site of Bazel-Sluis were selected to undergo microwear analysis. Specimen selection was influenced by the size of the specimens, as the larger pieces did not fit under the metallographic microscope. No chemical or mechanical alteration traces were observed on tool surfaces prior to analysis.

The tools were analysed following an established method of combined ‘low-power’ (1-50x magnification) and ‘high-power’ (50-1000x magnification) approaches. Three microscope systems were used to analyse the microwear traces: an Olympus SX7 stereomicroscope was used (8x-56x magnification), where micrographs were taken by an Olympus SC100 camera and processed with Olympus Stream Basic 1.9.4 software. A Zeiss AxioZoom V16 motorised microscope system with up to 112x digital magnification was also used. Micrographs were taken with a Zeiss AxioCam 305 camera and processed with ZEN Core 3.4 software. And an Olympus BX53 metallographic microscope (50-200x magnification)with a Nikon D750 DSLR camera and Best Scientific 1.9X coupler. Here, micrographs were captured with Helicon Remote software and processed with Helicon Focus software. The tools were cleaned in an ultrasonic bath with Derquim soap and ultrapurified water prior to use-wear analysis. All lithic tools were regularly cleaned with 96% alcohol and/or lighter fluid during the examination. Edge damages were registered with both the stereo and the metallographic microscopes, while the metallographic microscope was used to describe polish attributes, such as texture, topography, brightness, distribution, directionality and location. Striation attributes such as width, depth and length were also noted.

### Optical microscopy residue analysis

All lithics examined were stored individually in plastic bags. Work surfaces and microscope specimen tables were cleaned with 70-96% alcohol before each examination. During examination, lithic tools were handled with powder-free nitrite gloves. For high-magnification microscopy, the tools were fixed on a plastic slab with Blu Tack which was covered with parafilm. Parafilm was cleaned between specimens and was changed when damage occurred.

Possible residues were first identified, described and documented *in situ* using a Zeiss AxioZoom V16 motorised microscope with up to 112x digital magnification. Physical characteristics including colour, texture, and lustre were described as well as their location and their integrity with the surface, and the presence of any microtraces at the same locations. Some residues were also examined under the Olympus BX53 metallographic microscope at 200-500x magnification.

A subset of the microscopically identified residues were extracted for further analyses. Four areas of visible residue were removed by scraping the residues with sterile Dry Needling needles onto clean weighing papers. These residues were transferred into Eppendorf tubes and weighed. The heaviest sample was sent to ETH -Zürich to undergo ^14^C analysis. Nine samples were extracted for a staining analysis by placing the tools into clean zip lock bags, adding ultrapurified water to cover the area with adherent residue, and placing the bag in an ultrasonic bath for 15 minutes. A few microscope slides were prepared per sample by pipetting out water from the plastic bags and placing 1-2 drops onto to glass slides. The samples were dyed with Picrosirius Red following the protocol by Stephenson^63^. The slides were analysed using a Zeiss ImagerA2M microscope with polarised light but without an analyser for polarisation. Sixteen further samples were stained in the same manner after prior cleaning for use-wear analysis.

### Proteomic analysis

#### Protein Extraction

Protein extraction was performed by adapting the suspension trapping protocol described by Engels *et al.*^64^ . For each of the ‘soak’ samples 600µl was aliquoted from the total volume into 1.5 mL Eppendorfs and vacuum dried. The ‘swab’ samples were incubated at 25 °C in Milli-Q water for 30 min while shaking at 750 rpm (Eppendorf Thermomixer comfort). The samples were centrifuged (Eppendorf Centrifuge 5417R) and, after removal of the swab, vacuum dried. Both the ‘swab’ and ‘soak’ samples were resuspended in extraction buffer ((5% SDS (Invitrogen, 15553-027) + 50mM TEAB (Sigma-Aldrich, 90360-100ML)). For the ‘microscope plate’ samples, the nail polish sealing the cover-glasses was removed with acetone (Sigma-Aldrich, 179124-1L), after which 23µl of SDS buffer was used to resuspend sample residues on both the glass plate and the cover glass. All three sample types subsequently underwent protein reduction by DTT (Chemlab, CL00.0481.0025), at a final concentration of 20.8mM for 30min at 37°C in the dark. Next, alkylating agent MMTS (Sigma-Aldrich, 64306-10ML) was added to a final concentration of 20mM for 10 min at room temperature in the dark. The denatured proteins were precipitated with phosphoric acid (Chemlab, CL00.0605.1000) at pH 1.

Proteins were trapped on HiPure Viral Mini columns (Magen Biotechnology, China; C13112) after adding 165µl binding/washing buffer (100mM TEAB in 90% Methanol; Chemlab, CL00.1377.1000). The columns were centrifuged for 30 sec (4000 rpm, 25°C) between each of the three washing steps to elute the binding/washing buffer, and the first elution was reloaded on the column to decrease protein loss. The proteins were digested on-filter with 0.5µg trypsin/Lys-C (ProMega, V5073) in 40 µL of 50mM TEAB at 47°C for 2h. The peptides were eluted from the column with 30µl of 50mM TEAB, followed by 30µl of 0.1% formic acid (FA) (Biosolve, 2324) and finally 30µl of 50% acetonitrile (Chemlab, CL00.0194.1000). All three elution steps were performed with 1 min incubation then 1 min centrifugation. The samples were vacuum-dried and resuspended in 0.1% FA for LC-MS/MS analysis.

#### Instrumental Measurement

The samples were analysed using a Waters Acquity M-Class UPLC system coupled to a Sciex ZenoTOF 7600 mass spectrometer in data-dependent acquisition mode. The peptides were trapped on a YMC Triart C18 guard column (3µm, 5×0.3mm) and separated on a YMC Triart C18 (3µm, 150×0.3mm) analytical column using an optimised non-linear 20 minute gradient of 1.5 to 36% solvent B (0.1% FA in acetonitrile) in solvent A (0.1% FA in water). For the microscope ‘slide’ samples a clean new column was used. The clean column was tested and primed on a K652 cell lysate.

Precursor scans were acquired for 0.1s over a mass range of 300-1600*m/z*. Up to 40 precursors with an intensity threshold of 150 counts per second, a dynamic exclusion of 6 seconds after 2 occurrences and a charge state between 2+ and 5+ were fragmented per cycle using collision-induced dissociation. The fragment spectra were acquired for 0.015s over a mass range of 100-2000*m/z*, resulting in a cycle time of 0.920s.

#### Data Analysis

Raw datafiles were peak-picked by the MSConvert^74^ peak-picking algorithm into MGF file format. The MGF files were submitted to Mascot Daemon (version 2.8.2, Matrix Science, London, UK), and searched as merged inputs with three different search strategies: 1) the manually curated CollagenDB^64^ (mammalian taxa only, 12299 sequences) and Universal Contaminants database^75^ (downloaded 03/2023, 381 sequences); 2) an *in silico*-digested Major Seed Storage Proteins database (9683 sequences), a Milk Proteins database (871 sequences), and a common mammalian muscle and plasma protein database (18817 sequences), plus the Universal Contaminants database; 3) the SwissProt database^76^ (downloaded 01/2025, 572970 sequences) and Universal Contaminants database. These searches are referred to as the ‘CollagenDB’, ‘CerealKiller’, and ‘SwissProt’ searches. The enzyme was set to semiTrypsin with a maximum of 1 missed cleavage. Methylthio (C, +45.987721 Da) was added as a fixed modification, and Deamidation (NQ, +0.984016 Da), and Oxidation (M, +15.994915 Da) were added as variable modifications. Oxidation (P, +15.994915 Da) was additionally used for the CollagenDB search, and Phospho (ST, +79.96633) for the CerealKiller search. The fragment error was set to 30 ppm and the peptide error tolerance was 10 ppm.

For the three whole-dataset searches, Mascot search results were restricted to a significance threshold of P<0.01, with sub-set proteins disabled, requiring two unique significant peptides per protein, for export to .csv files. Data was filtered by removing PSMs derived from the proteases used during sample preparation (*Sus scrofa* trypsin and *Pseudomonas aeruginosa* Lys-C/protease IV). Outputs from these three searches were then combined for analysis.

For species inference analysis in ClassiCOL, a merged file was searched per tool (including all extractions) in Mascot against the CollagenDB^64^ and Universal Contaminants database^75^, after which result files were extracted in .csv data format restricted to a significance threshold of p<0.01 with one significant peptide per protein and with sub-sets enabled. Mascot .csv output files were used as input for the ClassiCOL algorithm version 1.0.1. The ClassiCOL taxonomy parameter was restricted to Mammalia, as the Mascot results did not display collagen peptides of non-mammalian origin.

NCBI BLASTp searches were performed against the ClusteredNR database with no taxonomic restrictions.

Full Methods for the modern saliva extract can be found in Supplementary Information.

### Radiocarbon dating

Due to the very small mass of the residue (<10 mg C), the standard measurement involving graphitization of the carbon could not be applied. Instead the sample was directly measured on CO_2_ in the Gas Ion Source of the MICADAS at ETH -Zürich^77,78^. An elemental analyser was used to convert the sample to CO2. Calibration of the radiocarbon date was performed using the IntCal20 curve ^79^ in the OxCal online version 4.4.

## Supporting information

BLASTp searches plant peptides

## Data Availability

Raw and processed mass spectrometry proteomics data from samples, blanks, and instrument flushes, and study-specific database FASTA files, are available to download via the PRIDE partner repository with the dataset identifier PXD068258 and 10.6019/PXD068258. The authors declare that all other data supporting the findings of this study are available within the paper and its supplementary information files.

## Acknowledgements

This study was supported by the Special Research Fund (BOF) of Ghent University, under funding grants BOF.GOA.2022.0002.03 and BOF.GOA.2022.0002.02 awarded to the Regional Outlook on Ancient Migration (ROAM) project (A.B., É.H., I.D.G., P.C. and M.D.). Additional funding was provided by ProGenTomics (A.B., I.E., D.D., M.D.). Residue analyses (optical and histological) carried out at the DANTE Laboratory, Sapienza University of Rome, were supported by the ERC under the EU’s Horizon 2020 programme (GA No. 639286, HIDDEN FOODS).

## Contributions

A.B., I.E. and É.H. conceived and designed the study. I.E. extracted and measured lithic proteomics samples, R.S.D extracted and measured the modern saliva samples, and A.B. carried out proteomics data analysis. É.H. performed use-wear, optical microscopy analysis of residue and histological staining analysis. E.C. supervised and performed optical microscopic residue and histological staining analysis. P.C. acquired radiocarbon analysis. P.C. and M.D. supervised the study. A.B., É.H, I.E., P.C. and M.D. wrote and edited the manuscript. All authors provided feedback on the manuscript. I.D.G. and D.D. secured funding for the analytical work.

## Corresponding author

Correspondence to Alexandra Burnett and/or Maarten Dhaenens.

## Ethics Declarations

The saliva samples analysed in this study were collected with informed consent and form part of the Forensic Proteomics Biobank ‘Karakterisering van lichaamsmaterialen met behulp van op massaspectrometrie gebaseerde eiwitanalyse’ (ONZ-2023-0306). This biobank was approved by the Committee for Medical Ethics at Ghent University Hospital.

## Competing Interests

The authors declare no competing interests.

## Electronic Supplementary Information

- **PDF**: Supplementary Information: extended methods, results, discussion, figures & tables
- **TXTs**: BLAST searches for plant proteins

## Notes

### Competing Interest Statement

The authors have declared no competing interest.

### Summary of Updates

Similarities between sections of text in this manuscript pertaining to site information and certain methods, and previously published manuscripts, have been revised.

