## Supplementary Information for "Palaeoproteomics reveals bovine carcass processing with 6,000-year-old flint tools"

### Sample Overview

| *Study lithic code* | *Contact material* | *Motion of use* | *Visual residues* | *Residue placement* | *Staining* | *Proteomic sample code/s* | *Proteomic sample counts* | | | *ClassiCOL analysis* |
| --- | --- | --- | --- | --- | --- | --- | --- | --- | --- | --- |
|  |  |  |  |  |  |  | *Slide* | *Soak* | *Swab* |  |
| 4694 | Indeter. |  | White, fatty, amorph. | Mostly ridges & contact points | Dark red & greenish | 4694 5494 | 1 | 3 1 | 1 | *Bos taurus /primigenius* |
| 5258 | Indeter. |  | Dark brown, fatty, amorph. | Banded behind contact edge |  | 5258 |  | 9 | 1 | *Capra hircus* and/or *Capreolus capreolus* [inferred] |
| 7561 | H. animal | Dynamic, grinding | White, fatty, amorph., some light brown | Edge scars, background smears |  |  |  |  |  |  |
| 7634 | H. animal, meat/bone | Dynamic, longitudinal | White, fatty, fibrous | Mostly in edge cracks |  | 7634 |  | 1 | 1 | Pecora  [low conf.] |
| 7876 | N/A | N/A |  |  |  |  |  |  |  |  |
| 7961 | H. animal | Dynamic | White, clumped, dry | Concentrated in scars | orange | 761 7861 7961 | 1 | 1  11 |  | *Bos taurus /primigenius & Muntiacus* |
| 7976 | Dry hide | Transversal, scraping | White, beige, brown, dry | Edge retouch & scraping edge |  | 7976 |  | 1 | 1 | Pecora  [low conf.] |
| 7978 | H. animal | Dynamic, crushing | White, fatty amorph.; Anatidae barbules with fatty white tissue | Scars & background; across surface |  |  |  |  |  |  |
| 8001 | H. animal | Dynamic, crushing | White, fatty, amorph. | Scars, some areas bubbly | Ruby red, orange | 6408 | 1 |  |  | *Bos taurus /primigenius*  [low conf.] |
| 8013 | Med. h. animal | Dynamic | White, fatty, amorph.; Anatidae barbules | Scars, background |  | 8013 | 1 | 1 | 1 | *Bos taurus /primigenius* |
| 8062 | H. animal | Dynamic | White, light grey, fatty amorph. | Diffuse | Dark red | 8062 |  | 1 | 1 | Pecora  [low conf.] |
| 9232 | Med. h. animal | Dynamic | Plant fibres | No clear connection to use-wear |  |  |  |  |  |  |
| 10068 | Indeter. animal |  | White, fatty amorph. | Edge scars, surface smears |  | 10068 | 1 | 2 |  | Pecora  [low conf.] |
| 10935 | Fresh hide | Scraping, transversal | White, fatty amorph. | Concentrated in scars |  | 10935 |  | 1 | 1 | Ambiguous  [low conf.] |
| 11064 | H. animal | Dynamic |  |  |  | 11064 |  | 1 | 1 | Ambiguous  [low conf.] |
| 11208 | Hard stone | Diagonal | Black, dark brown crystals | Embedded in scars |  |  |  |  |  |  |
| 11686 | H. animal | Dynamic |  |  | Orange, gold |  |  |  |  |  |
| 11821 | Bone/antler | Dynamic |  |  |  |  |  |  |  |  |
| 13552 | Soft animal | Dynamic |  |  |  |  |  |  |  |  |
| 13887 | Hide | Scraping, transversal | White, amorph. clumps; beige, brown spots | Edge scars; edge scars & background |  | 13887 |  | 1 |  | Bovinae  [low conf.] |
| 15202 | Med. h. animal | Dynamic | Beige, brown, black, greasy clumps | Cracks | Dark red | 15202 |  | 1 |  | Pecora  [low conf.] |
| 16261 | Med. h. animal | Dynamic | Beige, white, greasy, black spots | Scars |  |  |  |  |  |  |
| 16970 | H. animal | Dynamic | Beige, red, black fatty amorph. | Thick and thin films | Orange, gold | 16970 |  | 1 |  | Failed |
| 17664 | Uninterpretable | N/A |  |  |  |  |  |  |  |  |
| 17665 | Indeter. |  | White, fatty amorph.; fibres | Thick layer; across large areas | Orange & green | 17665 | 2 | 1 |  | Bovinae  [low conf.] |
| 17679 | H. animal | Non-specific | White, fatty, amorph. | Thickest at deep scar near edge | Orange |  |  |  |  |  |
| 17697 | H. animal | Dynamic |  |  |  |  |  |  |  |  |
| 18407 | Med. h. animal | Scraping, transversal | White, brown, fatty amorph; bright white crystals | Edge & non-edge scars | Ruby red, orange | 18407 |  | 1 |  | Failed |
| 18423 | Hide | Transversal | Bright white crystal with brown, black matter | Cracks |  | 18423 18623 | 1 | 4 |  | *Bos taurus*  */primigenius*  [low conf.] |
| 18532 | H. animal | Dynamic | White, beige, orange, brown fatty amorph. | Mostly connected to scars | Orange | 18532 |  | 1 |  | Failed |
| 18556 | Fresh hide | Scraping, transversal | White, beige, fatty amorph. | Scars, some background |  | 18556 |  | 1 |  | Failed |

Table S1: Overview of the analyses performed for all tools included in the study. ‘Indeter.’ = ‘Indeterminate’; ‘H.’ = ‘hard’; ‘Med.’ = ‘medium’; ‘amorph.’ = ‘amorphous’; ‘Low conf.’ = sample failed to meet the scoring thresholds outlined in Engels et al. 2025; ‘Failed’ = no classification was possible.

### Use-wear Analysis

| *Lithic code* | *UZ number* | *Degree of wear* | *Contact material main* | *Contact material sub1* | *Contact material sub2* | *Motion main* | *Motion sub* |
| --- | --- | --- | --- | --- | --- | --- | --- |
| 4694 | 1 |  | indeterminate |  |  |  |  |
| 5258 | 1 | probably used | indeterminate |  |  |  |  |
| 5258 | 2 | heavy | non-specific traces | abrasive | abrasive |  |  |
| 7561 | 1 |  | animal | unspecific | animal hard | dynamic | grinding |
| 7634 | 1 | heavy | animal | unspecific | animal hard | dynamic |  |
| 7634 | 2 |  | animal | meat | meat/bone | longitudinal | butchering |
| 7961 | 1 |  | animal | unspecific | animal hard | dynamic |  |
| 7976 | 1 |  | animal | hide | hide dry | transversal | scraping |
| 7978 | 1 |  | animal | unspecific | animal hard | dynamic | crushing |
| 8001 | 1 |  | animal | unspecific | animal hard | dynamic | crushing |
| 8013 | 1 | unsure | animal | unspecific | animal medium | dynamic | unspecific |
| 8062 | 1 | medium | animal | unspecific | animal hard | dynamic |  |
| 9232 | 1 | unsure | animal | unspecific | animal medium | dynamic | unspecific |
| 10068 | 1 | light | indeterminate |  |  |  |  |
| 10935 | 1 | medium | animal | hide | hide fresh | transversal | scraping |
| 11064 | 1 | medium | animal | unspecific | animal hard | dynamic |  |
| 11208 | 1 | heavy | inorganic | stone | stone hard | diagonal | unspecific |
| 11686 | 1 | medium | animal | unspecific | animal hard | dynamic | unspecific |
| 11821 | 1 | heavy | animal | unspecific | animal hard | dynamic |  |
| 13552 | 1 | medium | animal | unspecific | animal soft | dynamic |  |
| 13887 | 1 |  | animal | hide | hide unspecific | transversal | scraping |
| 15202 | 1 | heavy | animal | unspecific | animal medium | dynamic | unspecific |
| 16261 | 1 | medium | animal | unspecific | animal medium | dynamic |  |
| 16970 | 1 |  | animal | unspecific | animal hard | dynamic |  |
| 17664 | 0 | no traces |  |  |  |  |  |
| 17665 | 1 | light | indeterminate |  |  |  |  |
| 17679 | 1 | light | animal | unspecific | unspecific |  |  |
| 17697 | 1 |  | animal | unspecific | animal hard | dynamic |  |
| 18407 | 1 | medium | animal | unspecific | animal medium | transversal | scraping |
| 18423 | 1 | medium | animal | hide | hide unspecific | transversal | unspecific |
| 18532 | 1 | unsure | animal | unspecific | animal hard | dynamic |  |
| 18556 | 1 | medium | animal | hide | hide fresh | transversal | scraping |

Table S2: Summary of all interpreted used zones.

### Optical Microscopy Analysis of Residues

From the brownish appearance of some visible collagenous fatty residues, it was surmised that a very rapid and forceful motion may have caused slight burning of the matter through friction between the tool and hard animal matter, probably bone. This is suggested in part because of a similar pattern observed on experimental lithic tools, while the distribution of the discoloured residues and the colour pattern made burning by proximity of the tools to fires seem unlikely. During bone processing, possibly because of the heat generated by the motions, brown residues can sometimes appear. Through examination of both experimental and archaeological tools, it is observed that bone-working activities are often associated with striations that appear slightly brown and may show signs of burning. In our opinion, this may be linked to the heat produced during actions of tool use. The oxidation of organic residues such as blood and bone marrow, which are rich in iron, may occur under exposure to air or friction and this could also lead to brown or reddish discolorations. We have also observed that residues from worked bones on experimental tool surfaces often change colour, becoming yellow and glossy due to heat, and also display characteristic cracking. Due to the observation of the same phenomenon on experimental tools, discolouration by agents in the burial environment is also not thought to be responsible.

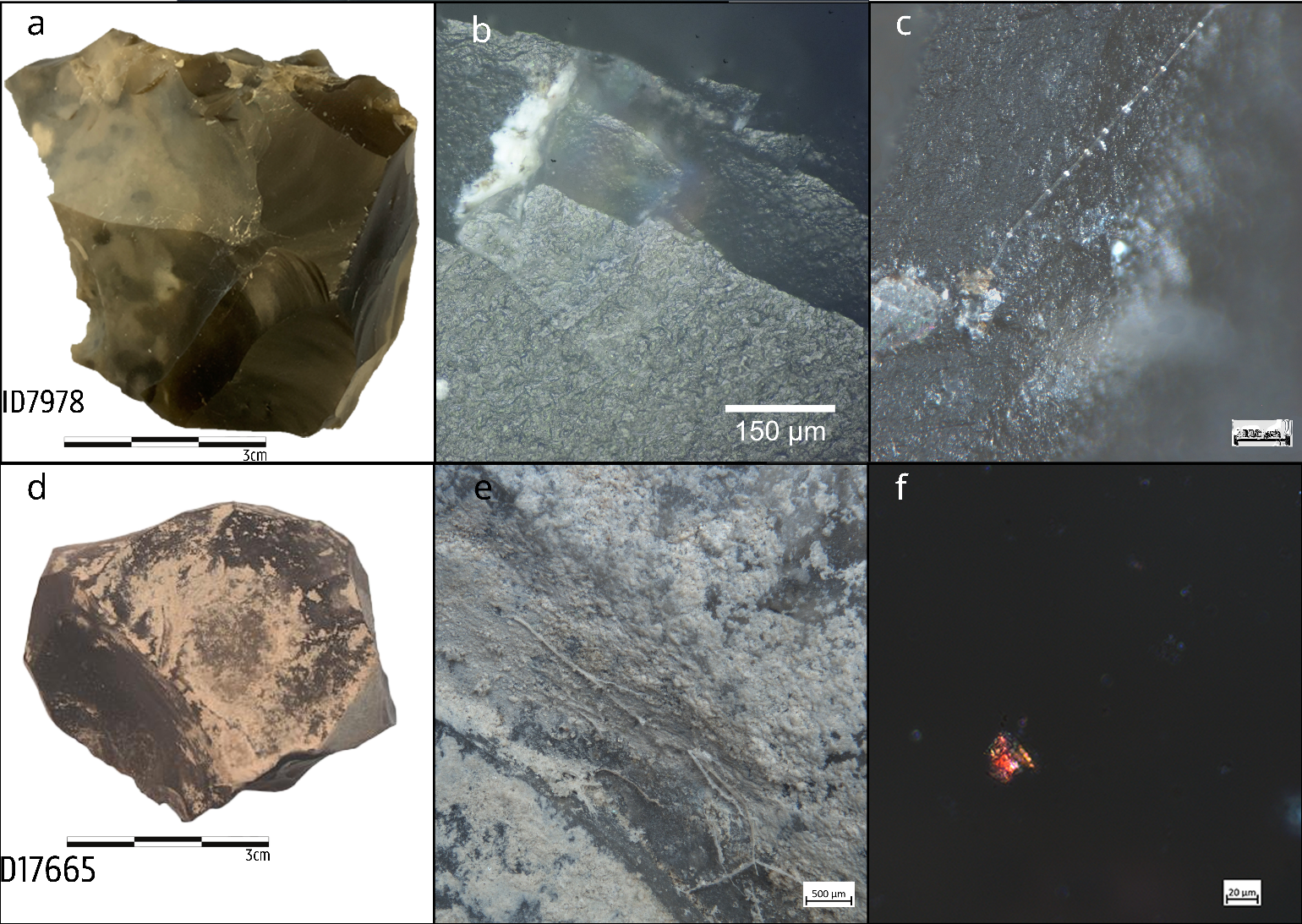

Figure S1: Faceted tools photographed with their visual residues and use-wear traces: a) tool 7978; b) traces of hard animal matter processed in a dynamic motion with smooth, flat, greasy, and bright polish with mixed directionality connected to edge scars with rounded edges; c) feather barbule of Anatidae family smashed onto the surface of 7978; d) tool 17665; e) thick layer of white amorphous matter with fibres; f) keratin residue from 17665 stained with Picrosirius red.

### Proteomic Analysis

#### Supplementary Methods

##### Database search strategies

The three databases used to provide identifications for the data in this study were selected for the following reasons. The SwissProt database contains all reviewed protein sequences from across the tree of life, and is commonly used as a starting point for samples with unknown protein content and for samples which contain protein from numerous (unknown) species. However, there is significant bias towards the identification of proteins from well-studied and model organisms, which are more likely to have reliable genomic, transcriptomic and proteomic evidence that contributes to their reliability and reviewed status. The CollagenDB was selected because it is the most comprehensive collagen database for proteomics, targeting one of the proteins that is most resistant to degradation and which can readily be used in a taxonomic assessment. The somewhat more miscellaneous ‘CerealKiller’ database was used for the following reasons: similarly to the rationale for the use of the CollagenDB, plant seed storage proteins are by nature resistant to degradation and were thought to be the most likely plant proteins to survive in detectable numbers. A group of soft tissue and muscle proteins was selected based on proteins identified on instruments analysed as part of forensic investigations, implemented here due to anticipated similarities with proteins deposited on tool surfaces through butchery.

It is possible that a plant database with more varied protein groups and more limited taxonomy (e.g. only from the 15 families identified) could produce results more closely aligned to prior palynological and macrobotanical findings from the site, but this would be at the risk of using an overly large database (with all the problems that entails) or, if using only reviewed sequences, including heavy bias towards model or heavily-studied organisms, including *Arabidopsis thaliana*.

The decision to merge all result files for each of the three main database searches was done largely to ensure that consistent *p*-values were applied across each dataset, given the variabilities between the sample types and the number of samples analysed per tool. This was intended to ensure faithful comparability between datasets, extraction types, and lithic tools.

##### Saliva Preparation

A 12mL saliva sample from a female donor was collected in 6x 2mL protein Lo-Bind Eppendorf tubes. Collection was undertaken at least 30 minutes after the last consumption of food or drink. Saliva samples were centrifuged for 10 minutes at 14,000 *g* and the resulting supernatants were combined in a 15mL falcon tube. A saliva dilution series: 1:2, 1:10, 1:100, 1:1000 and 1:10,000, was prepared using ultrapure water with 3 replicates per dilution point. Excess aliquots of saliva were vacuum-dried and stored at -20°C.

The suspension trapping protocol was carried out per the ProtiFi manufacturer’s protocol for the micro spin columns for 1-100µg of protein, with the modification that the reducing agent used was dithiothreitol (DTT; Chemlab, CL00.0481.0025) (ProtiFi, 2022). Briefly, samples were solubilised in 23µL SDS lysis buffer (5% SDS (Invitrogen, 15553-027) + 50mM TEAB (Sigma-Aldrich, 90360-100ML)). Samples were vortexed, sonicated for 5 minutes (Elma® Transsonic T460) and vortexed again. Reduction was performed by adding 1µL 137.5mM DTT and incubating at 37°C for 30 minutes at 750 rpm. Alkylation was performed by adding 1µL 500mM MMTS in isopropanol and incubating the samples in the dark for 10 minutes at 25°C. A protein crash was initiated by adding 2.5µL of 27.5% phosphoric acid to pH <1, and vortexing. 165µL of binding/wash buffer (100mM TEAB in 90% MeOH) was added to the samples which were then loaded onto the S-Trap column and centrifuged for 30 seconds at 4000 *g*. 150µL binding/wash buffer was loaded onto the column and centrifuged for 30 seconds at 4000 *g*, and repeated three times. The columns were transferred to clean 1.5mL protein Lo-Bind Eppendorf tubes. 1µg of Trypsin/Lys-C (ProMega, V5073) in 20µL 50mM TEAB was loaded on top of the filter and the samples were digested for 16 hours at 37°C. Peptides were eluted in three stages; (1) 30µL 50mM TEABC, (2) 30µL 0.1% FA and (3) 30µL of 50% ACN. The columns were centrifuged for 1 minute at 4000 *g* following the addition of each elution buffer and the resulting eluent vacuum-dried and stored at -20°C prior to resuspension in 20µL 0.1% FA for LC-MS/MS analysis.

2µL was injected from each saliva sample per analysis. Data was acquired on the instrumentation described in the main text. Peptides were separated on a linear gradient of 92.7 to 7.3% solvent A against solvent B (0.1% FA in water; 0.1% FA in ACN) over 21.5 minutes, at a flow rate of 5 µL/min. The ion source temperature was 200°C with a column temperature of 45°C and spray voltage of 4500V. Precursors of 400-1200 *m/z* were acquired in positive polarity mode. A maximum of 40 candidate ions with a charge state of 2^+^ to 4^+^ and whose intensity threshold exceeded 200 counts/sec were acquired. Candidate ions were excluded for 10 seconds after two occurrences. The precursor mass tolerance was ±100 ppm. MS/MS spectra of 140-1800 *m/z* were acquired. Ions were fragmented via CID with a de-clustering potential of 80 V, spray voltage of 4500 V and an accumulation time of 0.011 seconds.

All saliva datafiles were merged and submitted to Mascot Daemon using the ‘SwissProt’ search parameters described in the main text. Search results were restricted to the p<0.01 significance threshold with two unique significant peptides per protein before exporting to .csv files. Data was filtered by sorting peptide queries by descending score and removing duplicate query hits.

#### Supplementary Results

##### Database Searches

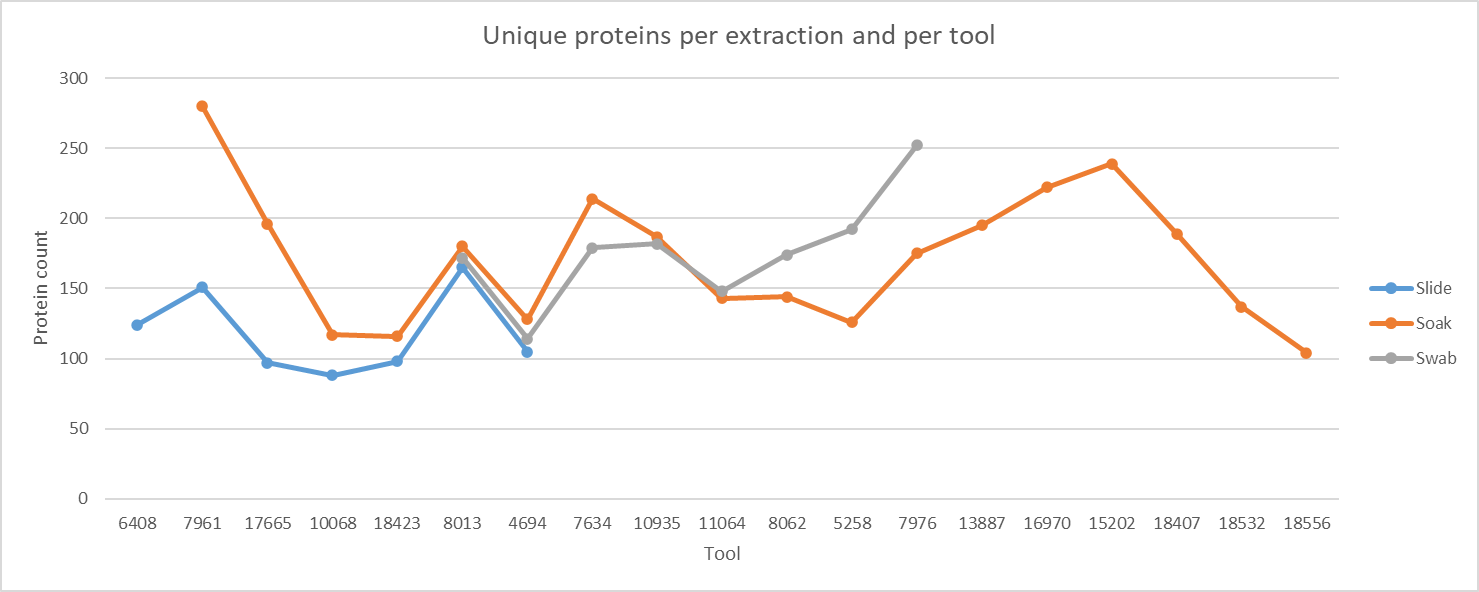

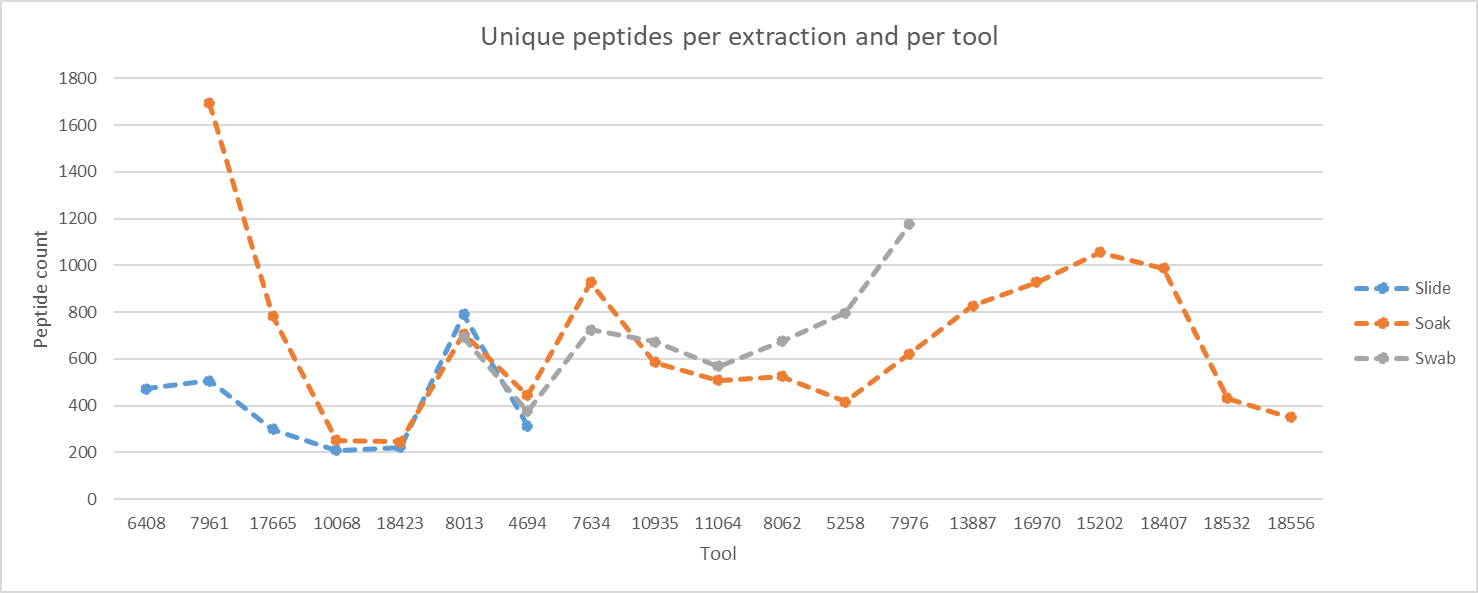

Figure S2: Variable protein and peptide content is seen per extraction and per tool.

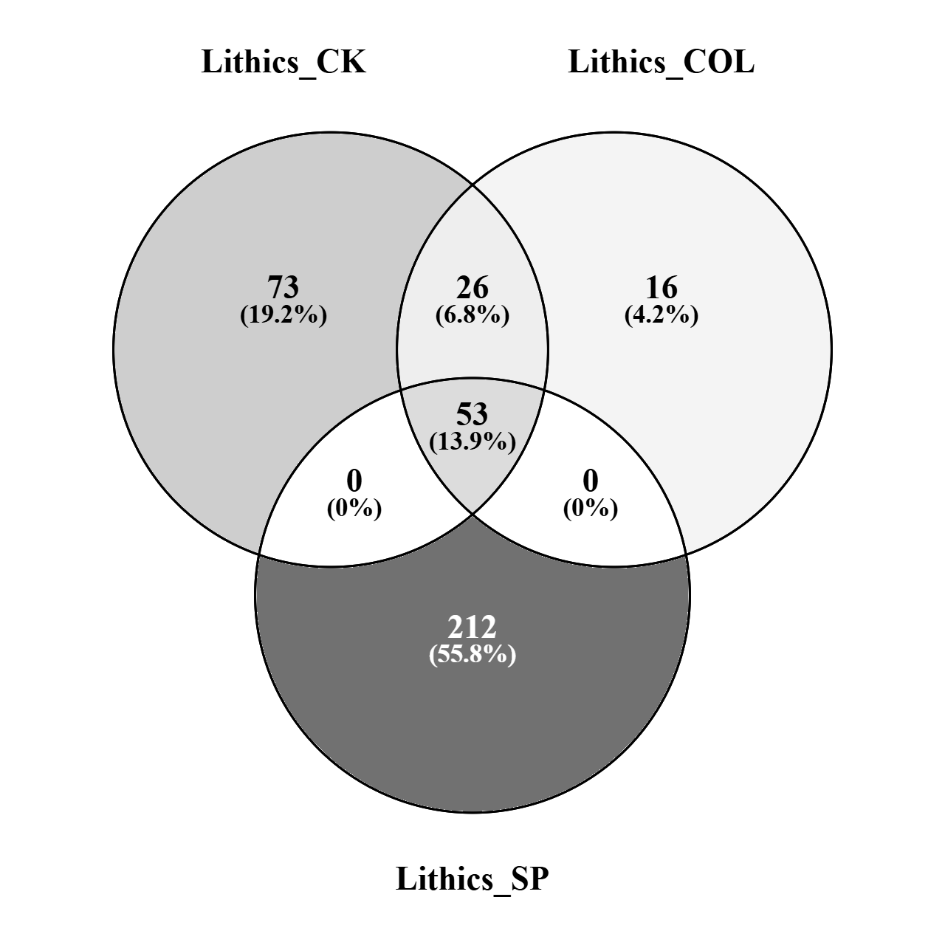

Figure S3: Overview of proteins and overlaps per search stratagem from all lithic samples.

‘CK’ = Cereal Killer search, ‘COL’ = Collagen search, ‘SP’ = SwissProt search. Comparison is made on the basis of unique protein descriptions. Some duplication may be present due to differing protein description titles in different databases (i.e. false unique-to-search entries, lower overlap counts). Diagram created using Venny 2.1.

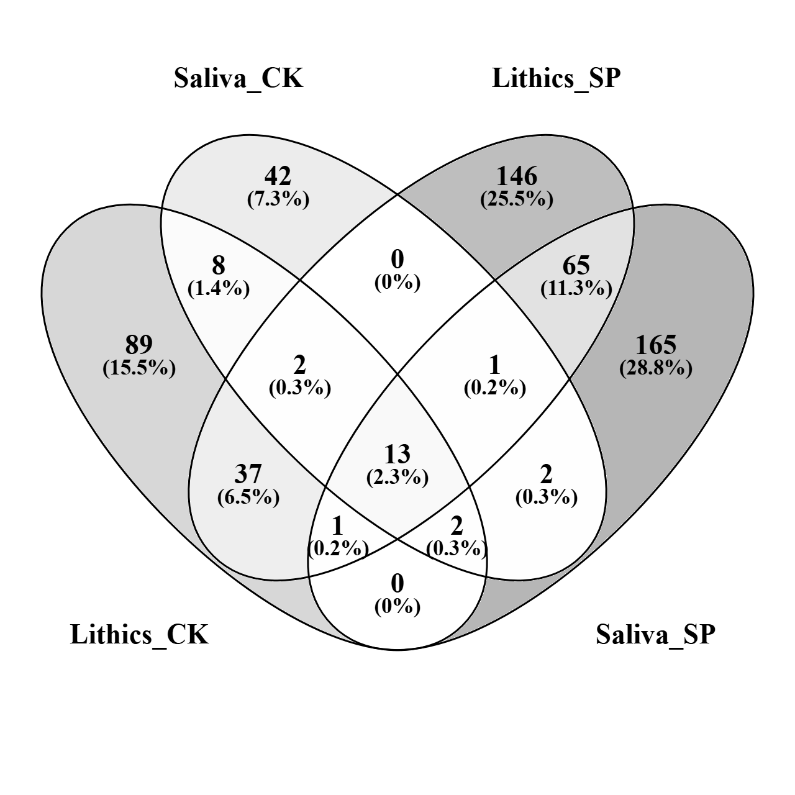

**13 universal common elements:**
14-3-3 protein zeta/delta OX=9913 GN=YWHAZ;
Filaggrin-2 OX=9606 GN=FLG2;
Keratin, type I cytoskeletal 10 OX=9606 GN=KRT10; Keratin, type I cytoskeletal 14 OX=9606 GN=KRT14; Keratin, type I cytoskeletal 16 OX=9606 GN=KRT16; Keratin, type I cytoskeletal 17 OX=9606 GN=KRT17; Keratin, type I cytoskeletal 9 OX=9606 GN=KRT9;
Keratin, type II cytoskeletal 1 OX=9606 GN=KRT1;
Keratin, type II cytoskeletal 1b OX=9606 GN=KRT77; Keratin, type II cytoskeletal 2 epidermal OX=9606 GN=KRT2;
Keratin, type II cytoskeletal 5 OX=9606 GN=KRT5;
Keratin, type II cytoskeletal 6A OX=9606 GN=KRT6A; Keratin, type II cytoskeletal 6B OX=9606 GN=KRT6B.

**92 common elements in "Lithics_CK-SP" and "Saliva_CK-SP"**:
14-3-3 protein zeta/delta OX=9913 GN=YWHAZ; Actin, cytoplasmic 1 OX=9913 GN=ACTB; Adenylyl cyclase-associated protein 1 OX=9913 GN=CAP1; Alpha-amylase OX=9913 GN=AMY2A; Alpha-enolase OX=9913 GN=ENO1; Chain A, Serotransferrin [Homo sapiens]; Chain X, LACTOTRANSFERRIN [Homo sapiens]; Elongation factor 1-alpha 1 OX=9913 GN=EEF1A1; Filaggrin-2 OX=9606 GN=FLG2; Fructose-bisphosphate aldolase A OS=Oryctolagus cuniculus OX=9986 GN=ALDOA; Gelsolin OX=9913 GN=GSN; Keratin, type I cytoskeletal 10 OX=9606 GN=KRT10; Keratin, type I cytoskeletal 12 OX=9606 GN=KRT12; Keratin, type I cytoskeletal 13 OX=9606 GN=KRT13; Keratin, type I cytoskeletal 14 OX=9606 GN=KRT14; Keratin, type I cytoskeletal 16 OX=9606 GN=KRT16; Keratin, type I cytoskeletal 17 OX=9606 GN=KRT17; Keratin, type I cytoskeletal 9 OX=9606 GN=KRT9; Keratin, type II cytoskeletal 1 OX=9606 GN=KRT1; Keratin, type II cytoskeletal 1b OX=9606 GN=KRT77; Keratin, type II cytoskeletal 2 epidermal OX=9606 GN=KRT2; Keratin, type II cytoskeletal 4 OX=9606 GN=KRT4; Keratin, type II cytoskeletal 5 OX=9606 GN=KRT5; Keratin, type II cytoskeletal 6A OX=9606 GN=KRT6A; Keratin, type II cytoskeletal 6B OX=9606 GN=KRT6B; Keratin, type II cytoskeletal 78 OX=9606 GN=KRT78; 6-phosphogluconate dehydrogenase, decarboxylating OX=9606 GN=PGD; Albumin OX=9606 GN=ALB; Alpha-1-antitrypsin OX=9606 GN=SERPINA1; Alpha-2-macroglobulin-like protein 1 OX=9606 GN=A2ML1; Alpha-amylase 1A OX=9606 GN=AMY1A; Alpha-enolase OX=9606 GN=ENO1; Annexin A1 OX=9606 GN=ANXA1; Antileukoproteinase OX=9606 GN=SLPI; Apolipoprotein A-I OX=9595 GN=APOA1; BPI fold-containing family B member 1 OX=9606 GN=BPIFB1; Calmodulin-like protein 3 OX=9606 GN=CALML3; Caspase-14 OX=9606 GN=CASP14; Catalase OX=9606 GN=CAT; Cathepsin D OX=9606 GN=CTSD; Cofilin-1 OX=9913 GN=CFL1; Complement C3 OX=9606 GN=C3; Cornulin OX=9606 GN=CRNN; Cystatin-A OX=9606 GN=CSTA; Cystatin-B OX=9595 GN=CSTB; Cystatin-S OX=9606 GN=CST4; Cystatin-SN OX=9606 GN=CST1; Desmocollin-2 OX=9606 GN=DSC2; Desmoglein-1 OX=9606 GN=DSG1; Fatty acid-binding protein 5 OX=9606 GN=FABP5; Fibrinogen gamma chain OX=9606 GN=FGG; Gelsolin OX=9606 GN=GSN; Glutathione S-transferase P OX=9606 GN=GSTP1; Glyceraldehyde-3-phosphate dehydrogenase OX=9606 GN=GAPDH; Haptoglobin OX=9606 GN=HP; IgGFc-binding protein OX=9606 GN=FCGBP; Immunoglobulin alpha-2 heavy chain OX=9606; Immunoglobulin gamma-1 heavy chain OX=9606; Immunoglobulin heavy constant alpha 1 OX=9606 GN=IGHA1; Immunoglobulin heavy constant gamma 3 OX=9606 GN=IGHG3; Immunoglobulin heavy constant mu OX=9606 GN=IGHM; Immunoglobulin J chain OX=9606 GN=JCHAIN; Immunoglobulin lambda constant 2 OX=9606 GN=IGLC2; Lactotransferrin OX=9606 GN=LTF; Lipocalin-1 OX=9606 GN=LCN1; L-lactate dehydrogenase A chain OX=9606 GN=LDHA; Lysozyme C OX=9595 GN=LYZ; Matrix metalloproteinase-9 OX=9606 GN=MMP9; Mucin-5B OX=9606 GN=MUC5B; Myeloperoxidase OX=9606 GN=MPO; Neutrophil defensin 1 OX=9606 GN=DEFA1; Neutrophil gelatinase-associated lipocalin OX=9606 GN=LCN2; Pancreatic adenocarcinoma up-regulated factor OX=9606 GN=ZG16B; Peptidyl-prolyl cis-trans isomerase A OX=9534 GN=PPIA; Peroxiredoxin-1 OX=9606 GN=PRDX1; Plastin-2 OX=9606 GN=LCP1; Polymeric immunoglobulin receptor OX=9606 GN=PIGR; Profilin-1 OX=9606 GN=PFN1; Prolactin-inducible protein OX=9606 GN=PIP; Protein S100-A7 OX=9606 GN=S100A7; Protein S100-A8 OX=9606 GN=S100A8; Protein S100-A9 OX=9606 GN=S100A9; Protein-glutamine gamma-glutamyltransferase E OX=9606 GN=TGM3; Pyruvate kinase PKM OX=9606 GN=PKM; Scavenger receptor cysteine-rich domain-containing protein DMBT1 OX=9606 GN=DMBT1; Serotransferrin OX=9606 GN=TF; Serpin B3 OX=9606 GN=SERPINB3; Thioredoxin OX=9606 GN=TXN; Triosephosphate isomerase OX=9606 GN=TPI1; Ubiquitin OX=9838; WAP four-disulfide core domain protein 2 OX=9606 GN=WFDC2; Zinc-alpha-2-glycoprotein OX=9606 GN=AZGP1.

Figure S4: Venn diagram showing overlap between lithic and saliva sample proteins.

‘CK’ = Cereal Killer search, ‘SP’ = SwissProt search. Taxa are identified by OS code: *Bos taurus* OX=9913; *Homo sapiens* OX=9606; *Oryctolagus cuniculus* OX=9986; *Gorilla gorilla gorilla* OX=9595; *Chlorocebus aethiops* OX=9534; *Camelus dromedarius* OX=9838. Counts are comprised of unique protein descriptions. Diagram created using Venny 2.1.

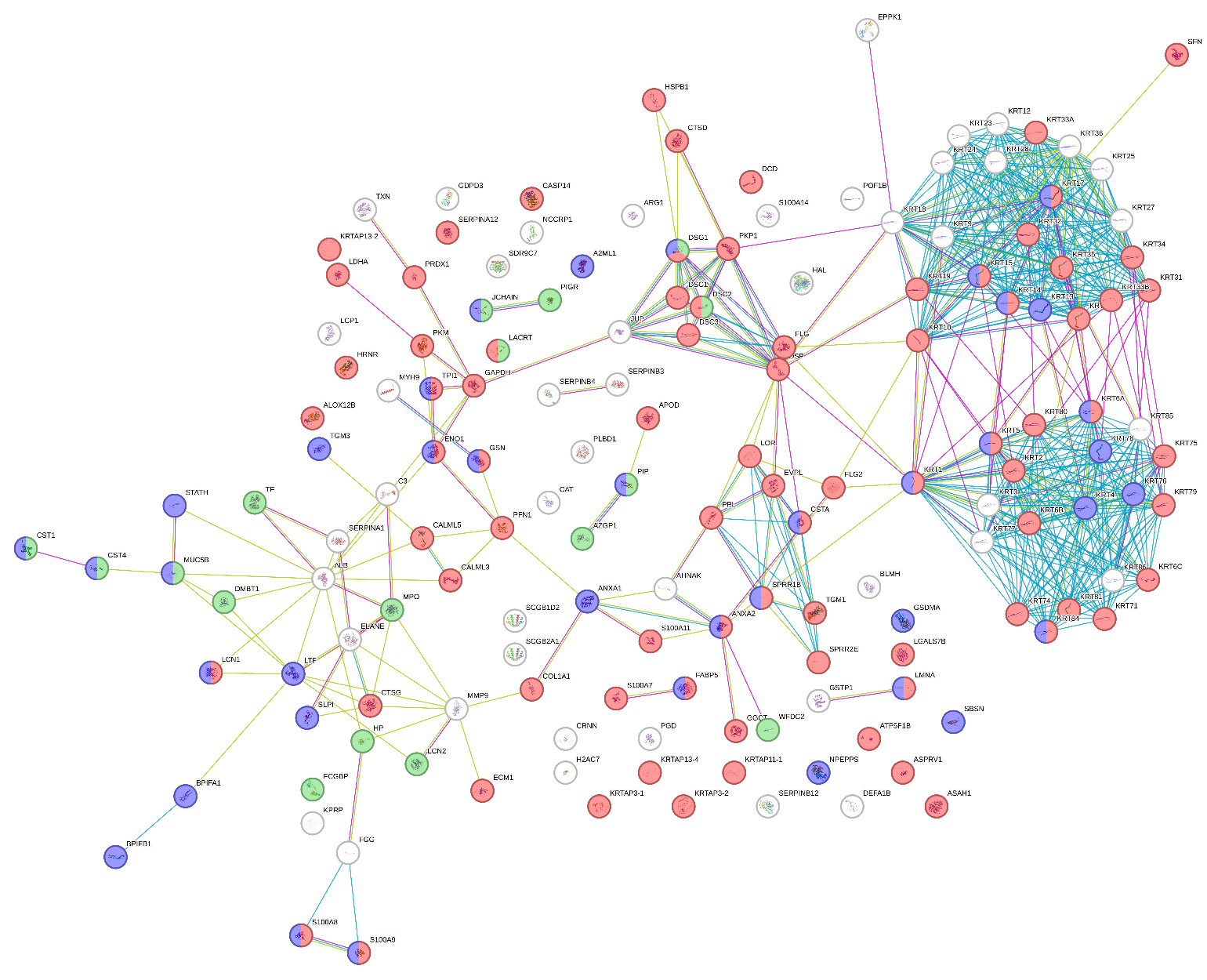

Figure S5: STRING mapping of human proteins demonstrates skin and salivary contamination.

All lithic tool proteins belonging to *Homo sapiens* were input to STRING and colour-coded by Tissue Expression (TISSUES) for ‘skin’ (red), ‘saliva’ (green), and ‘mouth’ (purple). Not all proteins were able to be mapped: the saliva biomarker alpha-amylase 1A and numerous immunoglobulins are absent from the figure. No clustering was applied; nodes are displayed as a physical subnetwork.
Permalink to STRING network: <https://version-12-0.string-db.org/cgi/network?networkId=bo2g0XVPTbY3>

| **PSMs per plant protein** |  |  |  |  |  |  |
| --- | --- | --- | --- | --- | --- | --- |
| ***Species*** (common name) | ‘CerealKiller’ search | | | ‘SwissProt’ search | | |
| Protein Description | **Slide** | **Soak** | **Swab** | **Slide** | **Soak** | **Swab** |
| ***Anacardium occidentale*** (cashew) | **18** | **80** |  |  | **10** |  |
| 11S globulin seed storage protein Ana o 2.0101 | 1 | 18 |  |  | 10 |  |
| 2S albumin | 17 | 62 |  |  |  |  |
| ***Arabidopsis thaliana*** (mouse-ear cress) | **5** | **43** | **14** |  |  |  |
| 12S storage protein CRB | 5 | 43 | 14 |  |  |  |
| ***Arachis hypogaea*** (peanut) | **3** | **22** | **6** |  |  |  |
| Chain A, Arachin Arah3 isoform | 2 | 11 | 2 |  |  |  |
| glycinin, partial | 1 | 11 | 4 |  |  |  |
| ***Avena sativa*** (oat) | **27** | **537** | **103** | **6** | **179** | **28** |
| 11S globulin | 9 | 175 | 36 |  |  |  |
| 12s globulin | 8 | 160 | 27 |  |  |  |
| 12S seed storage globulin 1 | 10 | 193 | 40 | 4 | 140 | 22 |
| avena alpha amylase trypsin inhibitor |  | 5 |  |  |  |  |
| Avenin |  | 4 |  | 2 | 39 | 6 |
| ***Carpinus fangiana*** (monkeytail hornbeam) |  | **18** | **3** |  |  |  |
| hypothetical protein FH972_002425 |  | 7 |  |  |  |  |
| hypothetical protein FH972_025376 |  | 11 | 3 |  |  |  |
| ***Carya illinoinensis*** (pecan) |  | **5** | **3** |  |  |  |
| hypothetical protein CIPAW_03G264300 |  | 5 | 3 |  |  |  |
| ***Cicer arietinum*** (chickpea) |  |  | **4** |  |  |  |
| provicilin-like |  |  | 4 |  |  |  |
| ***Cocos nucifera*** (coconut) |  | **6** |  |  |  |  |
| 11S globulin isoform 2 |  | 6 |  |  |  |  |
| ***Corylus avellana*** (European hazelnut) | **2** | **116** | **35** |  |  |  |
| 11S globulin seed storage protein Jug r 4-like | 2 | 116 | 35 |  |  |  |
| Chain A, 48-kDa glycoprotein |  | 6 |  |  |  |  |
| Cor a 16 precursor |  | 5 |  |  |  |  |
| ***Elaeis guineensis*** (African oil palm) |  | **2** |  |  |  |  |
| cocosin 1 isoform X2 |  | 2 |  |  |  |  |
| ***Ficus carica*** (fig) |  | **24** | **5** |  |  |  |
| hypothetical protein TIFTF001_040798 |  | 24 | 5 |  |  |  |
| ***Glycine max*** (soybean) | **1** | **4** | **21** |  | **4** | **22** |
| Beta-conglycinin, beta chain |  |  | 3 |  |  |  |
| Chain A, alpha prime subunit of beta-conglycinin |  |  | 3 |  |  |  |
| Chain A, GLYCININ G1 | 1 | 4 | 15 |  |  |  |
| Glycinin G1 |  |  |  |  | 3 | 16 |
| Glycinin G4 |  |  |  |  | 1 | 6 |
| ***Hevea brasiliensis*** (Pará rubber tree) |  | **9** | **1** |  |  |  |
| Rubber elongation factor protein |  | 9 | 1 |  |  |  |
| ***Juglans regia*** (English walnut) |  | **54** | **41** |  |  |  |
| 11S globulin seed storage protein 2-like |  | 22 | 16 |  |  |  |
| hypothetical protein F2P56_002973 |  | 6 | 8 |  |  |  |
| legumin B-like |  | 26 | 17 |  |  |  |
| ***Lolium rigidum*** (annual ryegrass) |  | **4** |  |  |  |  |
| 63 kDa globulin-like protein |  | 4 |  |  |  |  |
| ***Melastoma candidum*** (Asian melastome) |  | **3** |  |  |  |  |
| hypothetical protein MLD38_031864 |  | 3 |  |  |  |  |
| ***Musa acuminata*** (wild banana) |  |  |  |  | **20** | **8** |
| Glucan endo-1,3-beta-glucosidase |  |  |  |  | 20 | 8 |
| ***Oryza sativa Indica Group*** (indica rice) | **1** | **39** | **8** |  |  |  |
| hypothetical protein OsI_06562 | 1 | 39 | 8 |  |  |  |
| ***Oryza sativa Japonica Group*** (japonica rice) | **6** | **60** | **20** |  | **4** | **5** |
| glutelin | 6 | 60 | 20 |  |  |  |
| Glutelin type-A 1 |  |  |  |  | 4 | 5 |
| ***Parasponia andersonii*** (cannabiaceae family) | **3** | **8** |  |  |  |  |
| Bicupin, oxalate decarboxylase/oxidase | 3 | 8 |  |  |  |  |
| ***Phragmites australis*** (common reed) |  | **3** |  |  |  |  |
| glutelin type-B 5-like |  | 3 |  |  |  |  |
| ***Pisum sativum*** (garden pea) |  |  | **5** |  |  | **8** |
| Legumin A |  |  |  |  |  | 6 |
| Provicilin (Fragment) |  |  |  |  |  | 2 |
| Vicilin, partial |  |  | 5 |  |  |  |
| ***Prosopis alba*** (neltuma alba) |  | **2** |  |  |  |  |
| legumin type B-like |  | 2 |  |  |  |  |
| ***Prunus dulcis*** (almond) |  | **102** | **5** |  | **59** | **1** |
| Chain A, Prunin |  | 68 | 4 |  |  |  |
| Prunin 1 Pru du 6.0101 |  |  |  |  | 59 | 1 |
| RmlC-like cupins superfamily protein, partial |  | 34 | 1 |  |  |  |
| ***Sesamum angolense*** (Angolan sesame) |  | **12** |  |  |  |  |
| 11S globulin seed storage protein 2 |  | 12 |  |  |  |  |
| ***Sesamum indicum*** (sesame) |  | **4** | **2** |  | **3** |  |
| 11S globulin precursor isoform 4 |  | 4 | 2 |  |  |  |
| 11S globulin seed storage protein 2 |  |  |  |  | 3 |  |
| ***Solanum dulcamara*** (bittersweet) |  | **1** | **2** |  |  |  |
| 11S globulin seed storage protein Jug r 4-like |  | 1 | 2 |  |  |  |
| ***Theobroma cacao*** (cocoa) | **4** |  | **2** | **6** | **8** | **5** |
| 21 kDa seed protein |  |  |  | 4 | 8 | 5 |
| Vicilin OS=Theobroma cacao |  |  |  | 2 |  |  |
| vicilin, partial | 4 |  | 2 |  |  |  |
| ***Triticum aestivum*** (common wheat) | **3** | **14** | **1** |  |  |  |
| gamma-gliadin/LMW-glutenin chimera Ch2 precursor, partial | 3 | 10 | 1 |  |  |  |
| globulin 3 |  | 4 |  |  |  |  |
| ***Triticum turgidum subsp. durum*** (durum wheat) |  | **4** |  |  |  |  |
| unnamed protein product |  | 4 |  |  |  |  |
| ***Vicia sativa*** (common vetch) |  | **2** | **7** |  |  |  |
| legumin A |  | 2 | 7 |  |  |  |
| ***Zea mays*** (maize/corn) | **2** | **3** |  | **2** | **61** |  |
| 1,4-alpha-glucan-branching enzyme 2, chloroplastic/amyloplastic |  |  |  |  | 2 |  |
| Globulin-2 | 2 | 3 |  |  |  |  |
| Granule-bound starch synthase 1, chloroplastic/amyloplastic |  |  |  | 2 | 59 |  |
| ***Zizania latifolia*** (Manchurian wild rice) |  | **1** | **3** |  |  |  |
| glutelin precursor |  | 1 | 3 |  |  |  |
| **Extraction Total** | **75** | **1193** | **291** | **14** | **348** | **77** |

Table S3: Plant PSMs per protein.

The CollagenDB search was excluded from the table as it contained only 6 PSMs for a single species (the rubber tree, which is considered a common contaminant). Protein descriptions have been trimmed for clarity and duplicate descriptions from different DBs combined.

Figure S6: Bovine keratin coverage and unique peptides

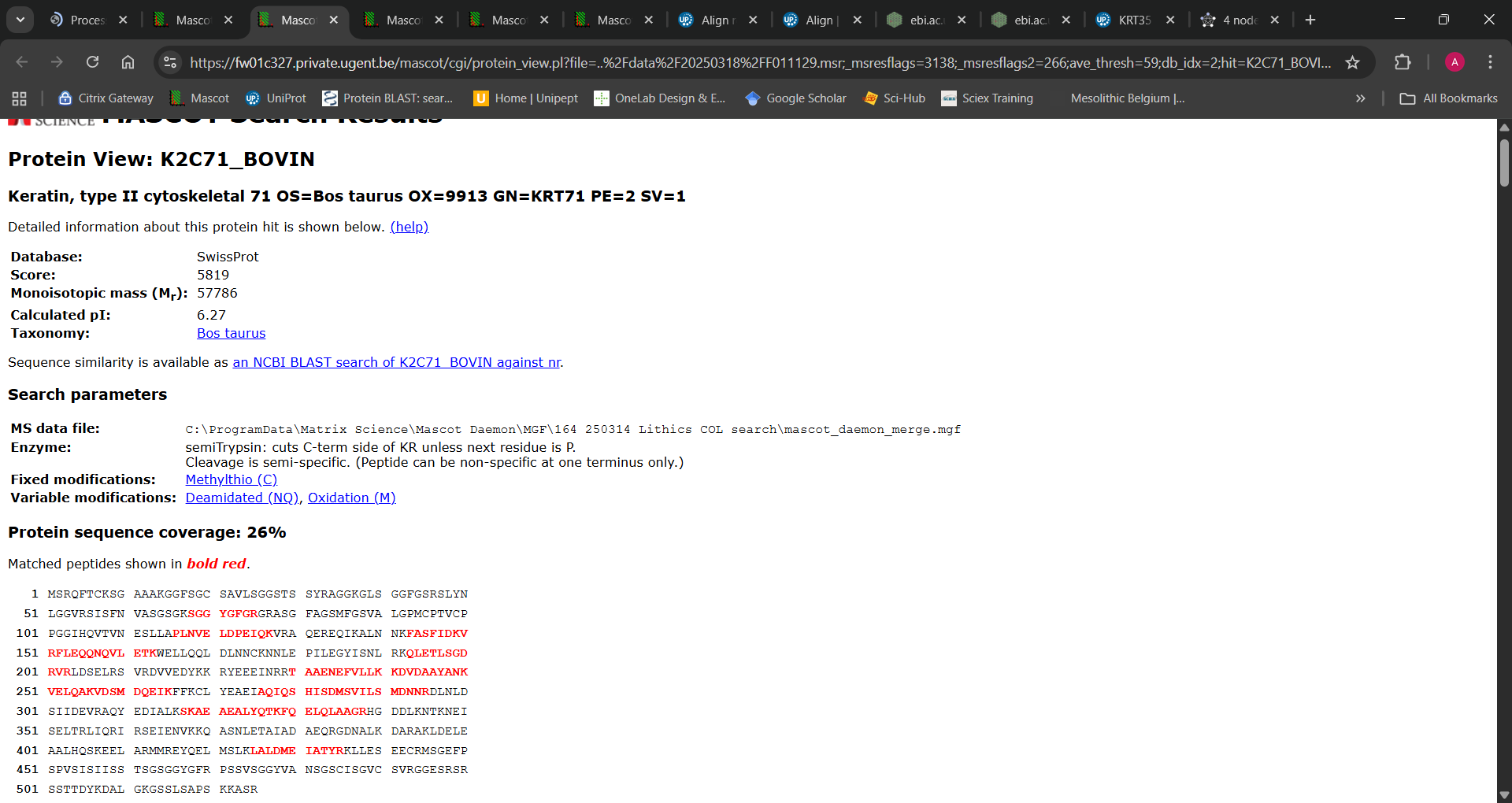

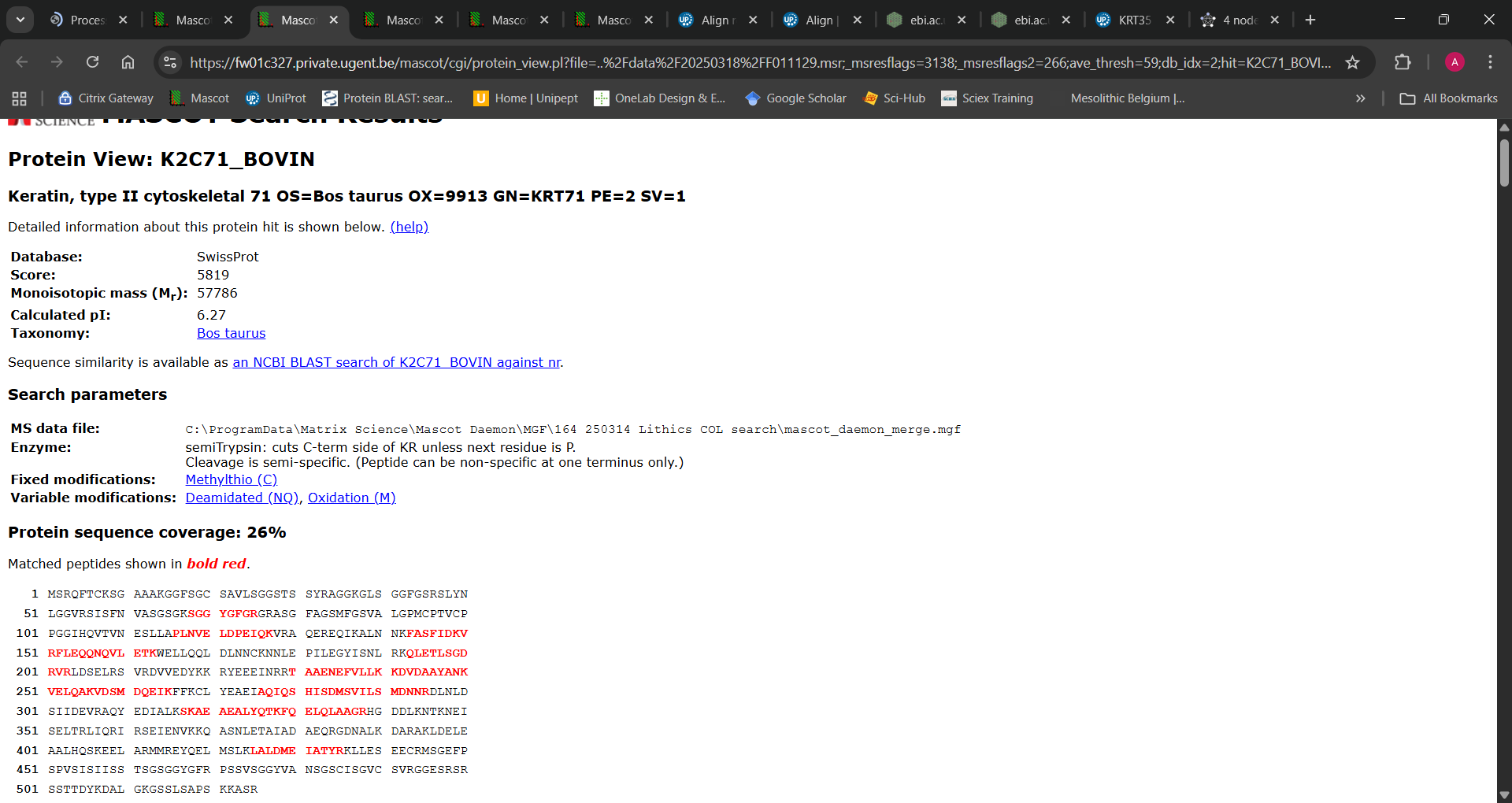

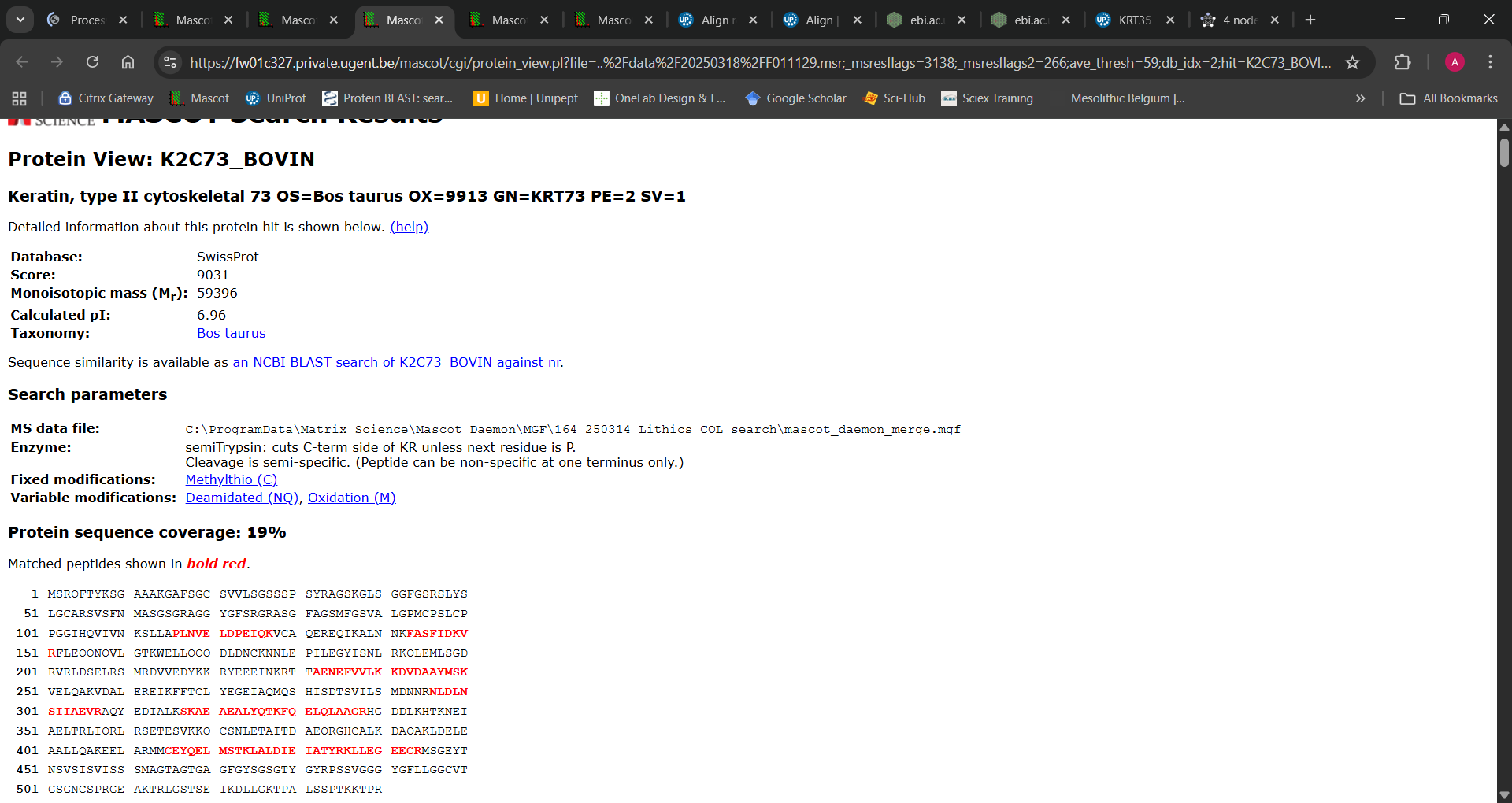

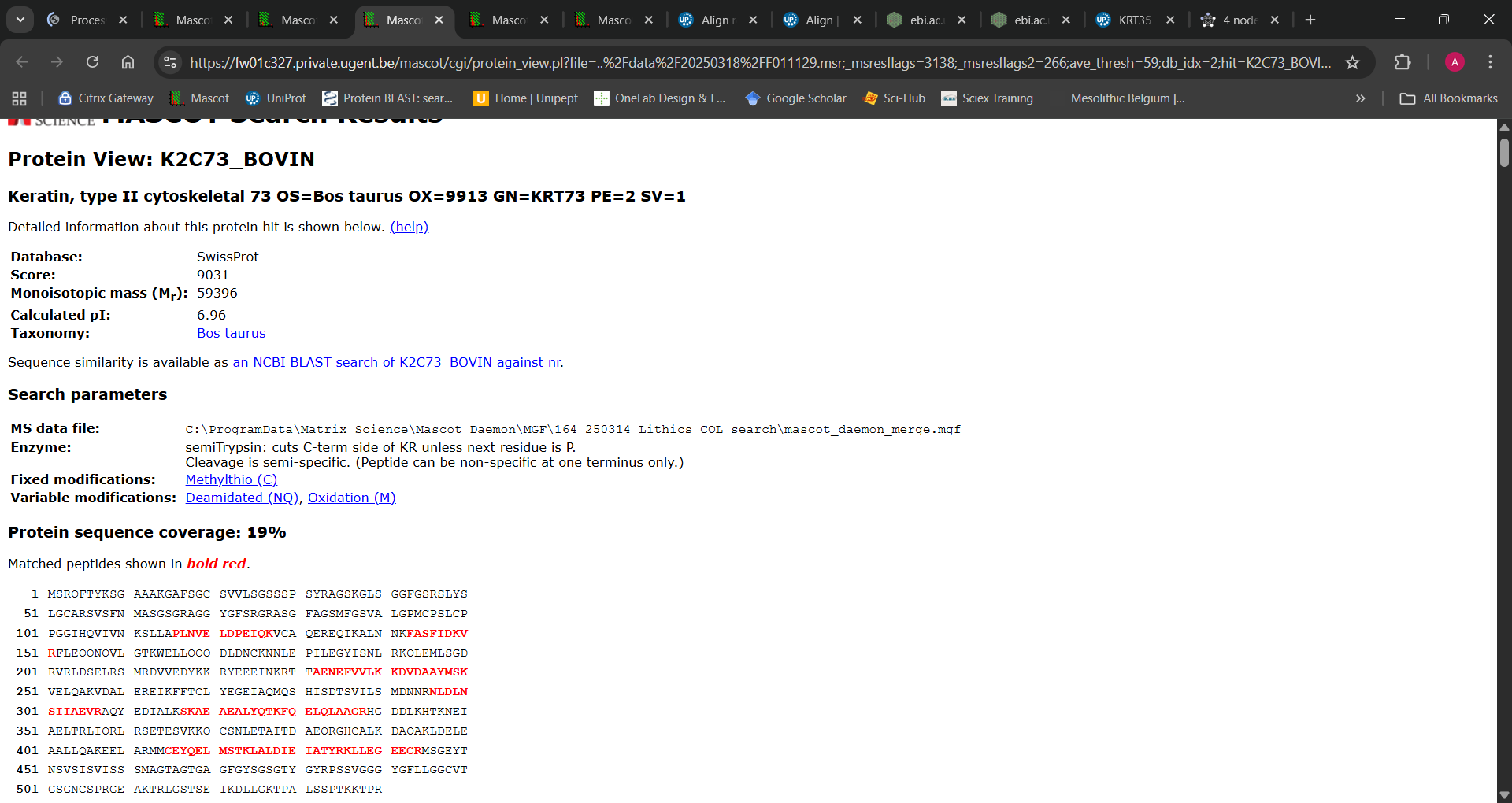

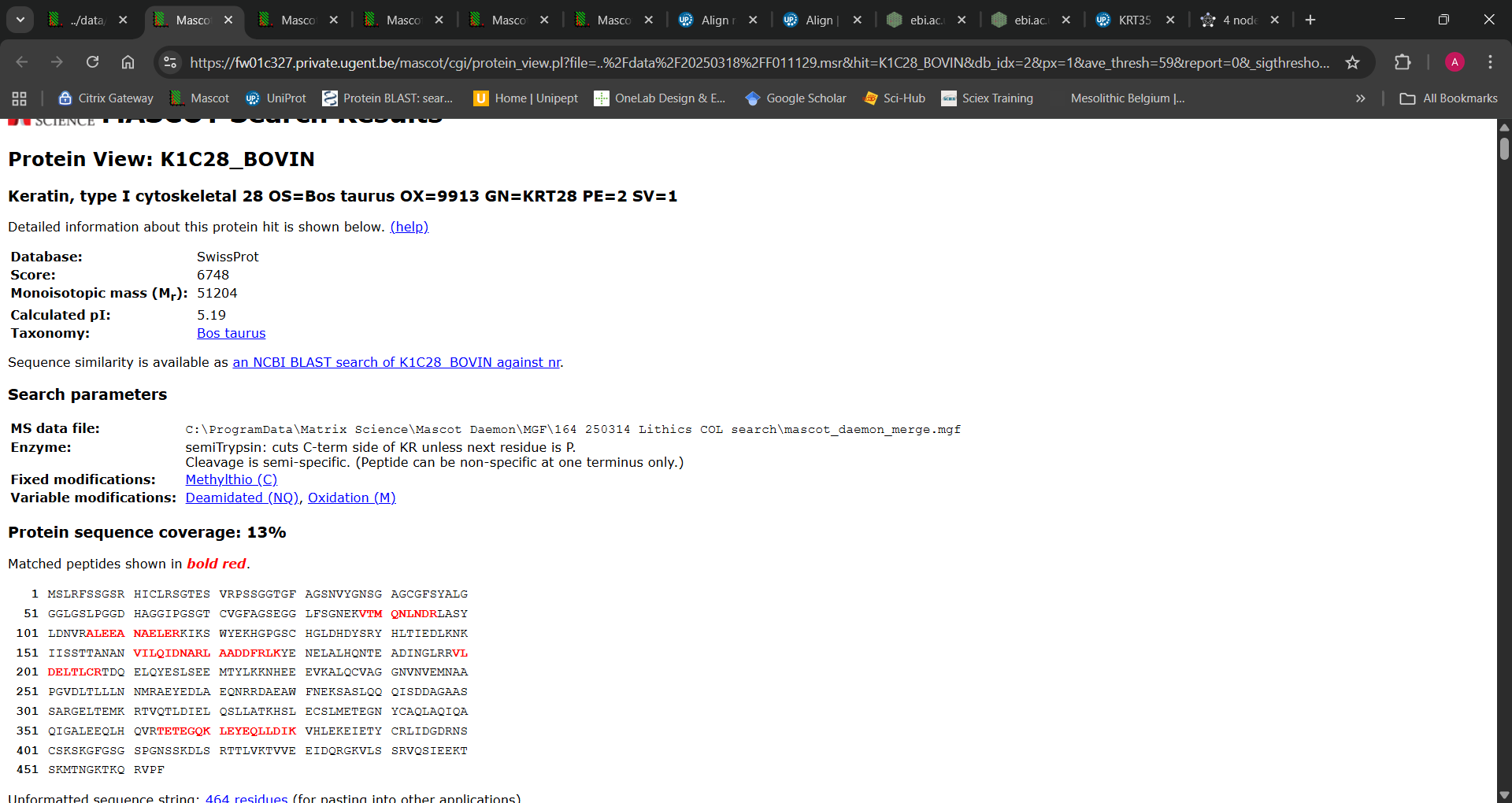

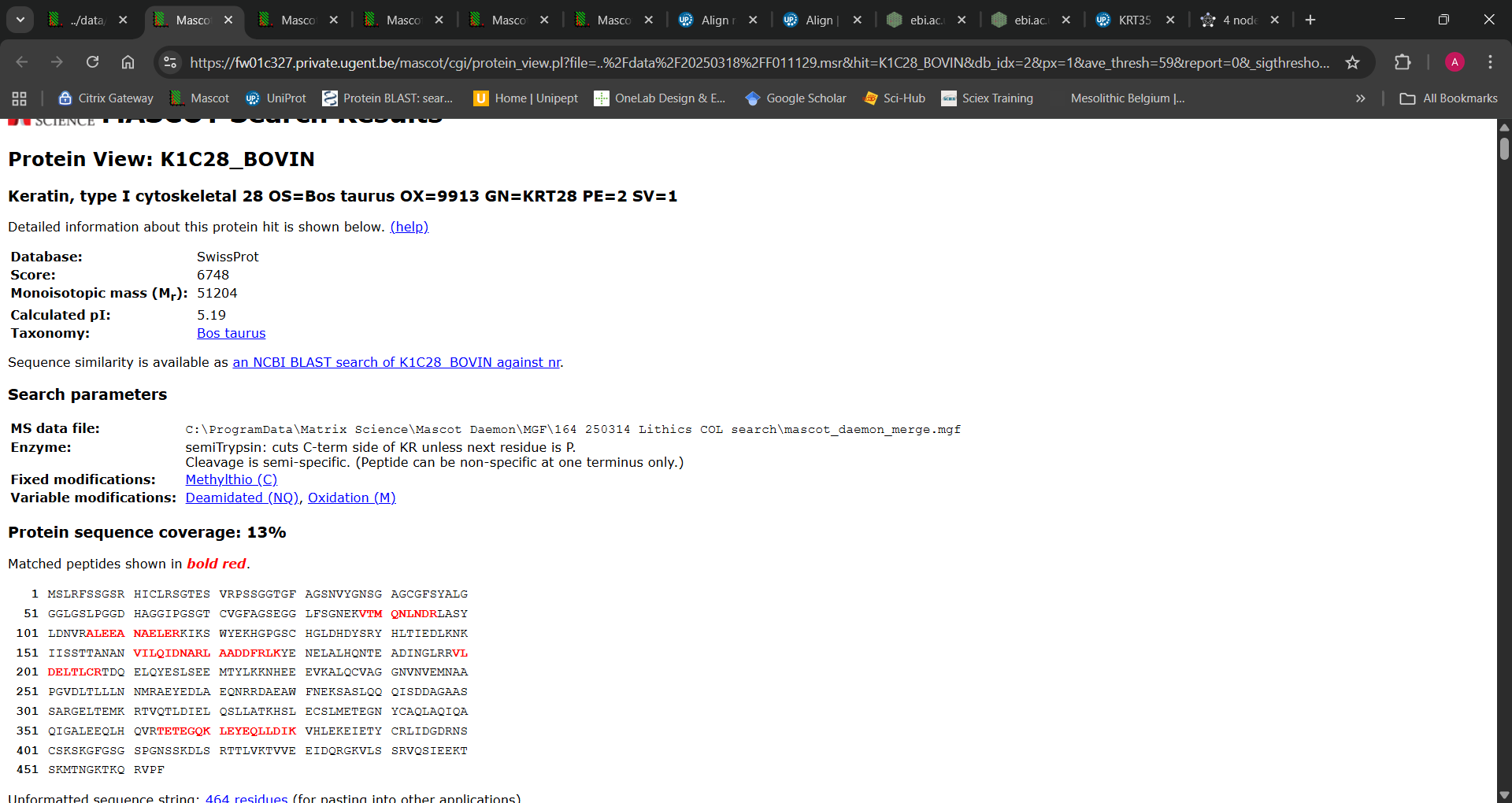

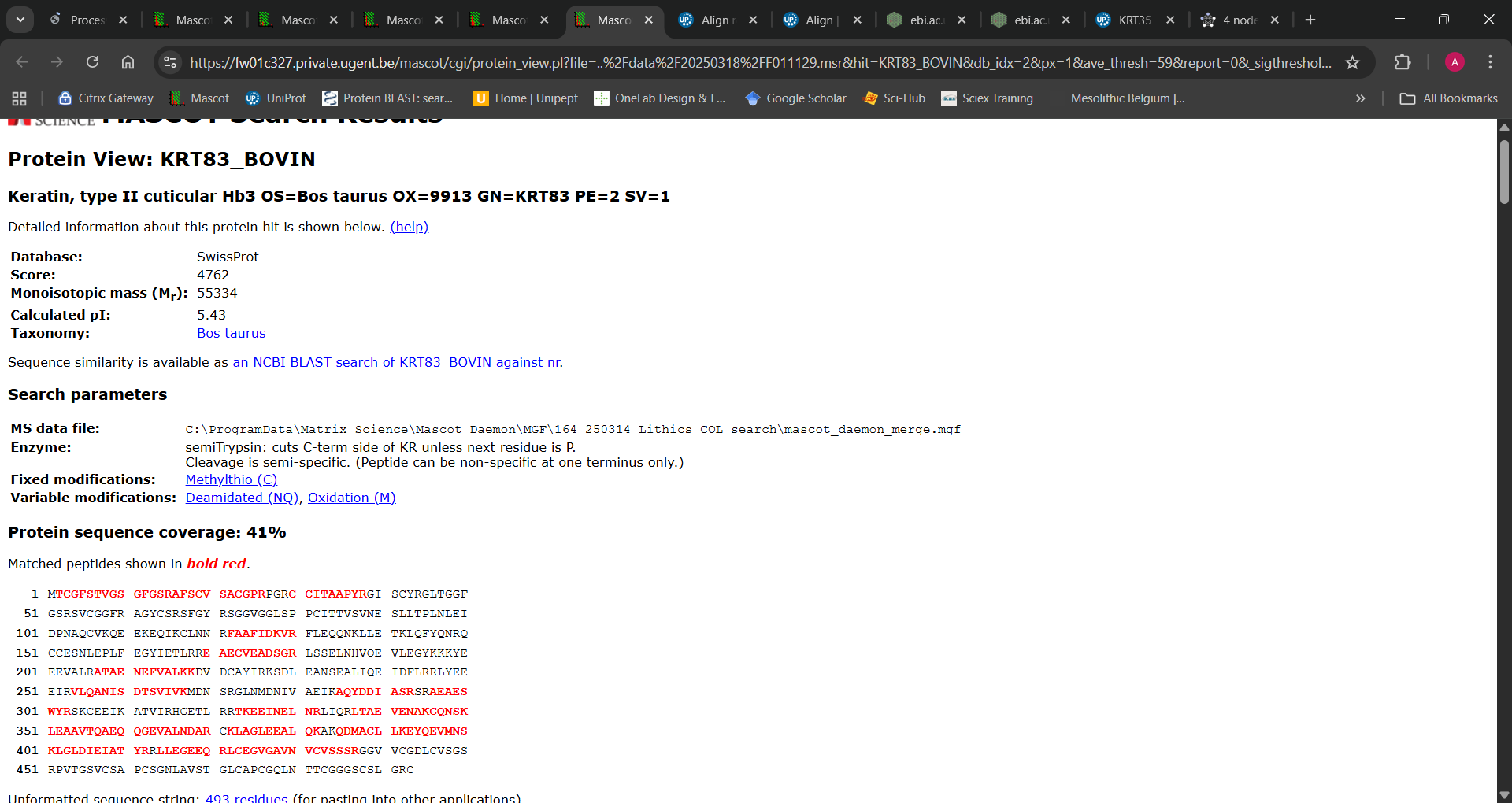

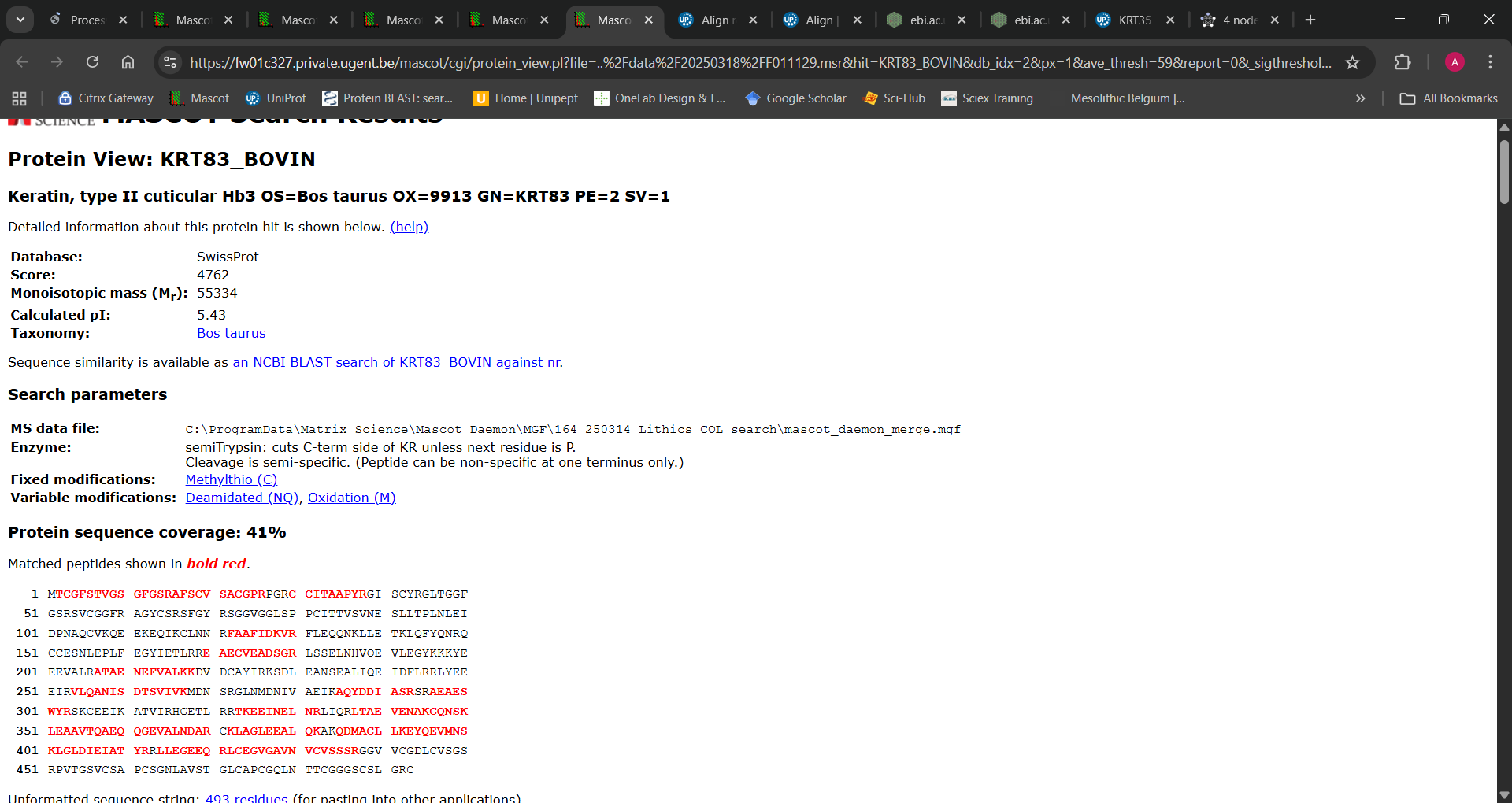

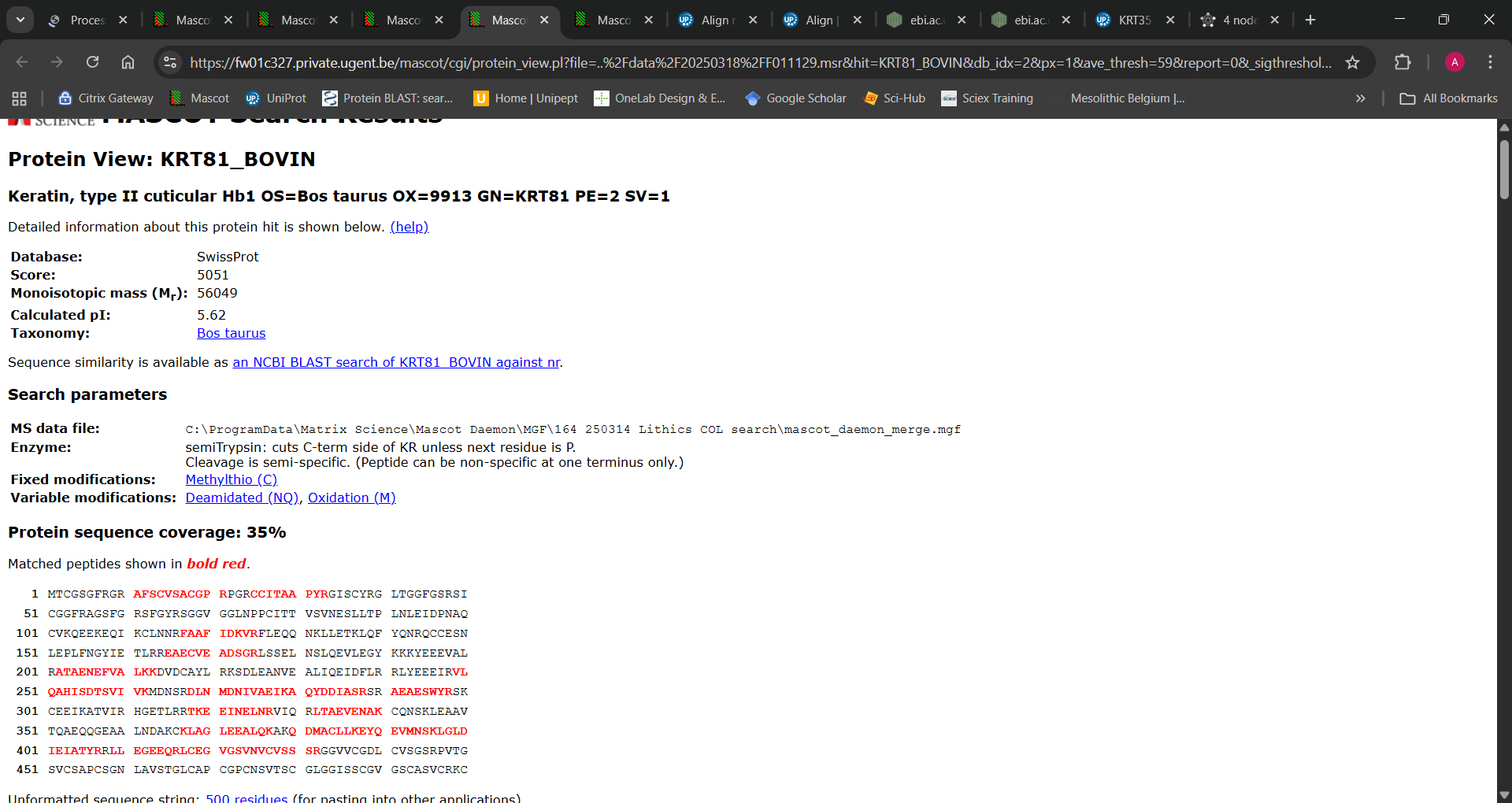

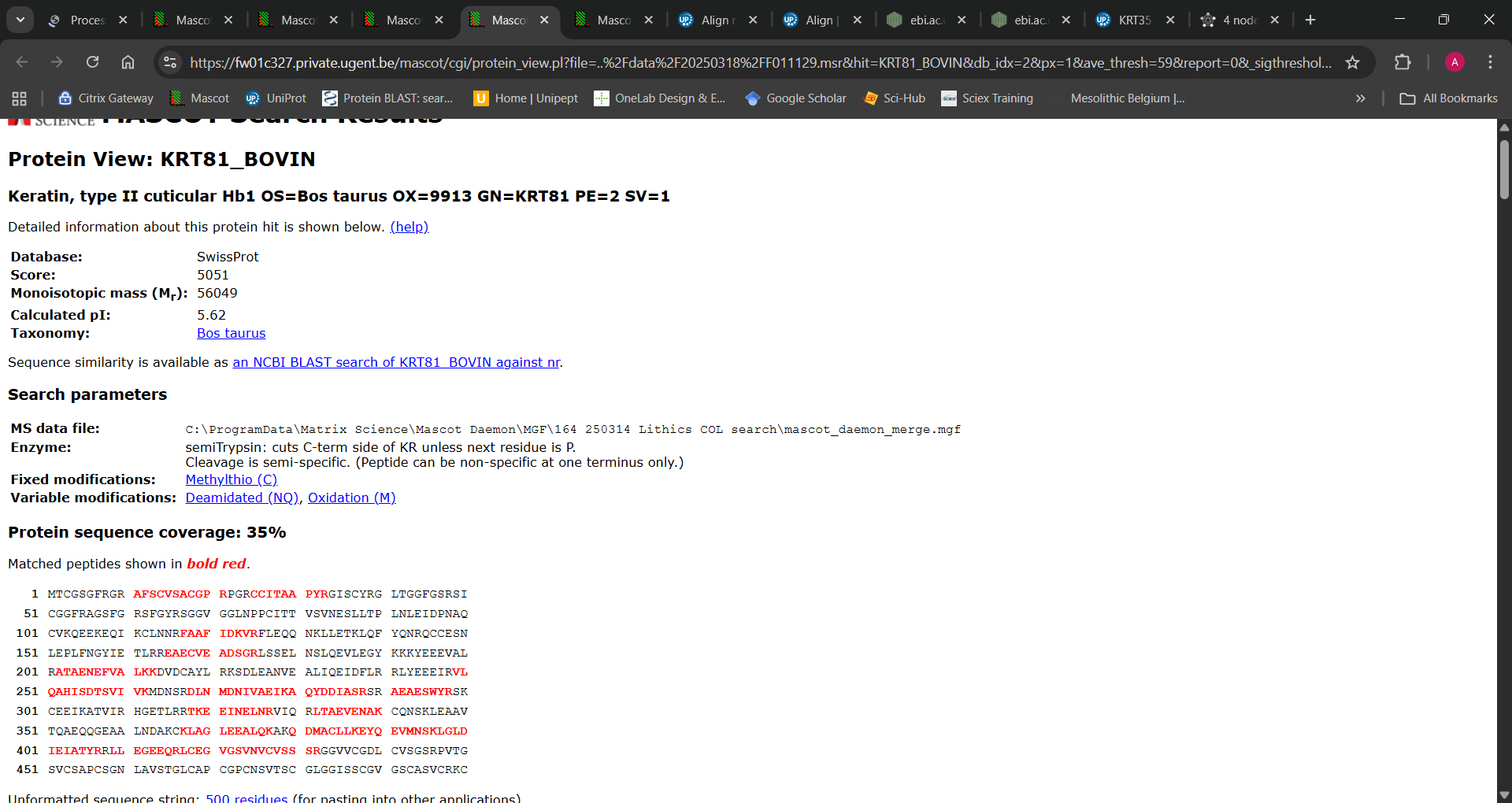

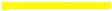

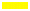

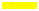

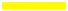

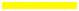

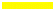

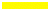

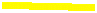

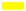

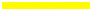

Peptides shown in red are identified in search results. Eight peptides highlighted in yellow are those which were not identified to any other protein in the dataset. Some peptides highlighted but not in red were identified in data outputs as ambiguous isobaric matches (e.g. DVDCAYIR -> DVDCAYLR).

| **PSMs per Protein (*Bos taurus*)** | **‘CerealKiller’ search** | | | | **CollagenDB search** | | | **‘SwissProt’ search** | | | | **Total** |
| --- | --- | --- | --- | --- | --- | --- | --- | --- | --- | --- | --- | --- |
| **Protein Description**  (OS=Bos taurus OX=9913) | **Slide** | **Soak** | | **Swab** | **Slide** | **Soak** | **Swab** | **Slide** | **Soak** | | **Swab** |  |
| 14-3-3 protein zeta/delta GN=YWHAZ |  | 49 | | 13 |  | 47 | 12 |  | 15 | | 6 | 142 |
| Actin, cytoplasmic 1 GN=ACTB | 31 | 495 | | 223 | 33 | 445 | 210 |  |  | |  | 1437 |
| Adenylyl cyclase-associated protein 1 GN=CAP1 |  |  | | 2 |  |  |  |  |  | |  | 2 |
| Albumin GN=ALB |  |  | |  | 15 | 306 | 109 | ***** | ***** | | ***** | 430 |
| Alpha-actinin-4 GN=ACTN4 |  |  | |  |  |  |  |  | 1 | | 3 | 4 |
| Alpha-amylase GN=AMY2A | 3 | 18 | | 6 | 3 | 14 | 3 | ***** | ***** | | ***** | 47 |
| Alpha-enolase GN=ENO1 |  | 37 | | 15 |  | 46 | 17 | ***** | ***** | | ***** | 115 |
| Alpha-S1-casein GN=CSN1S1 |  | 10 | | 3 |  | 7 | 3 |  |  | |  | 23 |
| Alpha-S2-casein GN=CSN1S2 | 8 | 72 | | 5 | 4 | 40 |  | 2 | 23 | |  | 154 |
| ATP synthase subunit alpha, mitochondrial GN=ATP5F1A |  |  | |  |  |  |  |  | 4 | | 4 | 8 |
| Beta-casein GN=CSN2 |  |  | |  | 8 | 22 | 10 | *Capra hircus* | | | | 40 |
| Beta-lactoglobulin GN=LGB | *Moschus berezovskii/Ovis aries* | | | | 4 | 32 | 5 | *Ovis aries musimon* | | | | 41 |
| Chloride intracellular channel protein 1 GN=CLIC1 |  |  | |  |  |  |  |  |  | | 3 | 3 |
| Clathrin heavy chain 1 GN=CLTC |  |  | |  |  |  |  |  |  | | 2 | 2 |
| Cofilin-1 GN=CFL1 |  |  | |  |  |  |  |  |  | | 4 | 4 |
| Collagen alpha-1(I) chain GN=COL1A1 | 52 | 16 | | 22 | 75 | 110 | 86 | 26 | 10 | | 21 | 418 |
| Collagen alpha-2(I) chain GN=COL1A2 | 30 | 2 | | 5 | 59 | 142 | 70 | 14 |  | |  | 322 |
| Elongation factor 1-alpha 1 GN=EEF1A1 | 3 | 41 | | 27 | 3 | 24 | 18 | Various taxa | | | | 116 |
| Endoplasmic reticulum chaperone BiP GN=HSPA5 |  |  | |  |  |  |  |  | 20 | | 19 | 39 |
| Gelsolin GN=GSN |  | 4 | | 4 |  | 3 | 3 |  | ***** | | ***** | 14 |
| Hemoglobin subunit alpha GN=HBA |  |  | |  |  | 24 |  |  | |  | | 24 |
| Kappa-casein GN=CSN3 | 11 | 37 | | 16 | 11 | 26 | 12 | 10 | 22 | | 12 | 157 |
| Keratin, type I cytoskeletal 28 GN=KRT28 | ***** | ***** | | ***** | ***** | ***** | ***** | 56 | 1022 | | 154 | 1232 |
| Keratin, type II cuticular Hb1 GN=KRT81 | ***** | ***** | | ***** | ***** | ***** | ***** | 25 | 467 | | 82 | 574 |
| Keratin, type II cuticular Hb3 GN=KRT83 |  |  | |  |  |  |  | 25 | 427 | | 81 | 533 |
| Keratin, type II cytoskeletal 71 GN=KRT71 | ***** | ***** | | ***** | *Mus musculus* | | | 30 | 268 | | 70 | 368 |
| Keratin, type II cytoskeletal 73 GN=KRT73 |  |  | |  | *Mus musculus* | | | 37 | 487 | | 178 | 702 |
| Large ribosomal subunit protein eL18 GN=RPL18 |  |  | |  |  |  |  |  |  | | 2 | 2 |
| Large ribosomal subunit protein P2 GN=RPLP2 |  |  | |  |  |  |  |  |  | | 5 | 5 |
| Plasma serine protease inhibitor GN=SERPINA5 |  | 2 | |  |  |  |  |  |  | |  | 2 |
| Protein disulfide-isomerase GN=P4HB |  | |  | |  |  |  |  |  | | 3 | 3 |
| Serine protease 1 GN=PRSS1 | 58 | 7 | | 1 | 56 | 8 | 3 | 32 |  | |  | 165 |
| Serotransferrin GN=TF |  |  | |  | 2 | 92 | 36 |  |  | |  | 130 |
| Small ribosomal subunit protein uS13 GN=RPS18 |  |  | |  |  |  |  |  | 1 | | 3 | 4 |
| Small ribosomal subunit protein uS2 GN=RPSA |  |  | |  |  |  |  |  | 7 | | 4 | 11 |
| Transitional endoplasmic reticulum ATPase GN=VCP |  |  | |  |  |  |  |  | 1 | | 6 | 7 |
| Tropomyosin beta chain GN=TPM2 |  |  | |  |  | 2 | 1 |  |  | |  | 3 |
| Tubulin alpha-1D chain GN=TUBA1D |  | 16 | | 23 |  | 8 | 24 |  |  | |  | 71 |
| Tubulin alpha-4A chain GN=TUBA4A |  | 16 | | 14 |  | 4 | 14 |  |  | |  | 48 |
| Tubulin beta-5 chain GN=TUBB5 |  | 2 | | 12 |  | 1 | 10 |  |  | | 3 | 30 |
| **Grand Total** | **196** | **824** | | **391** | **273** | **1403** | **646** | **257** | **2775** | | **667** | **7432** |

Table S4: All Bos taurus PSM counts identified in different extractions.
A red asterisk***** indicates the identification of an equivalent Homo sapiens protein for a given search. Alternative species names in red indicate equivalent protein identifications for the named species. Alternative taxon PSMs are not included in PSM totals.

| **PSMs per *Bos taurus* collagen** | **Protein** | |  |
| --- | --- | --- | --- |
| **Extraction**  Tool | **COL1A1** | **COL1A2** | **Total COL PSMs** |
| **Slide** | **75** | **59** | **134** |
| 4694 | 10 | 9 | 19 |
| 6408 | 12 | 13 | 25 |
| 7961 | 5 | 5 | 10 |
| 8013 | 14 | 10 | 24 |
| 10068 | 7 | 2 | 9 |
| 17665 | 19 | 14 | 33 |
| 18423 | 8 | 6 | 14 |
| **Soak** | **110** | **142** | **252** |
| 4694 | 10 | 17 | 27 |
| 5258 | 30 | 23 | 53 |
| 7634 |  | 4 | 4 |
| 7961 | 29 | 54 | 83 |
| 7976 | 1 | 2 | 3 |
| 8013 | 1 | 1 | 2 |
| 8062 | 1 | 2 | 3 |
| 10068 | 8 | 3 | 11 |
| 10935 | 1 | 2 | 3 |
| 11064 | 2 | 2 | 4 |
| 13887 | 3 | 10 | 13 |
| 15202 | 2 | 3 | 5 |
| 16970 |  | 1 | 1 |
| 17665 |  | 4 | 4 |
| 18407 |  | 1 | 1 |
| 18423 | 21 | 12 | 33 |
| 18532 | 1 | 1 | 2 |
| **Swab** | **83** | **70** | **153** |
| 4694 | 6 | 8 | 14 |
| 5258 | 15 | 13 | 28 |
| 7634 | 22 | 17 | 39 |
| 7976 | 3 | 2 | 5 |
| 8013 | 15 | 12 | 27 |
| 8062 | 9 | 6 | 15 |
| 10935 | 6 | 5 | 11 |
| 11064 | 7 | 7 | 14 |
| **Grand Total** | **268** | **271** | **542** |

Table S5: Bos taurus COL1A1 and COL1A2 PSMs per extraction type in all tools (Collagen search)

##### ClassiCOL outcomes

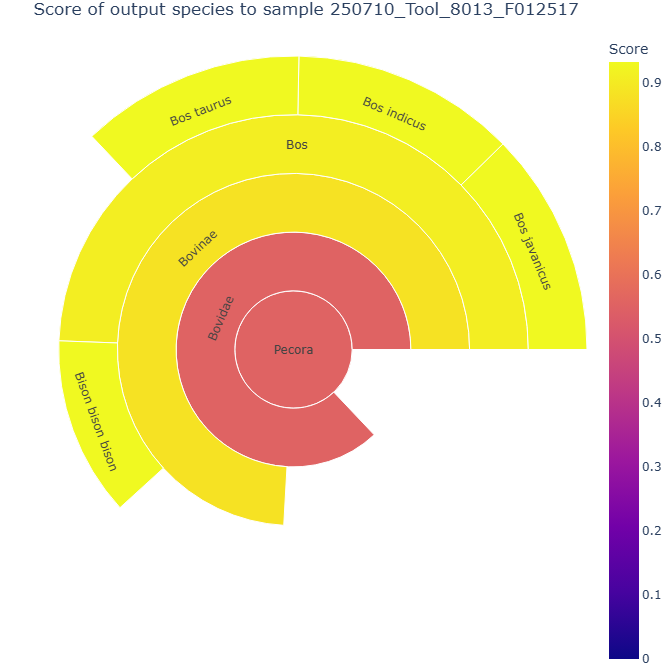

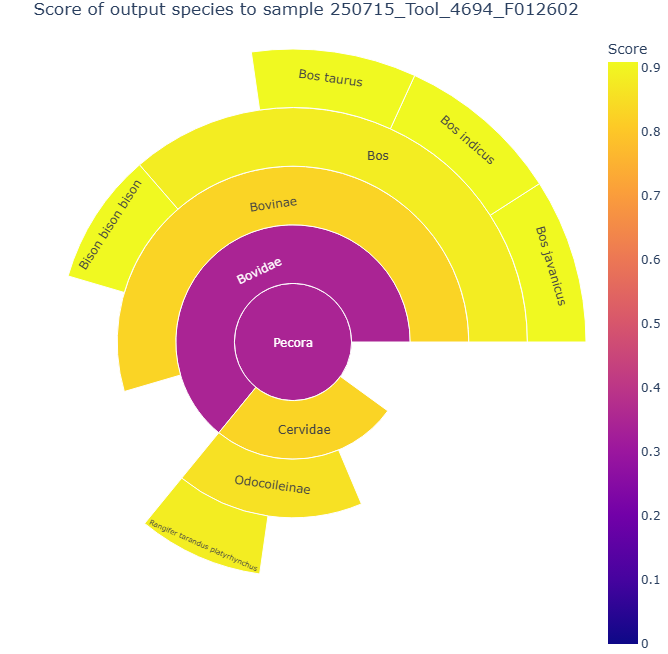

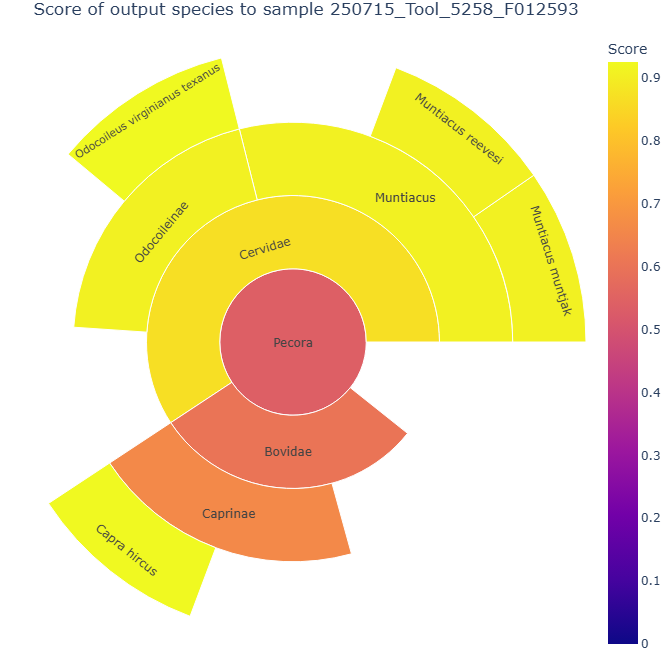

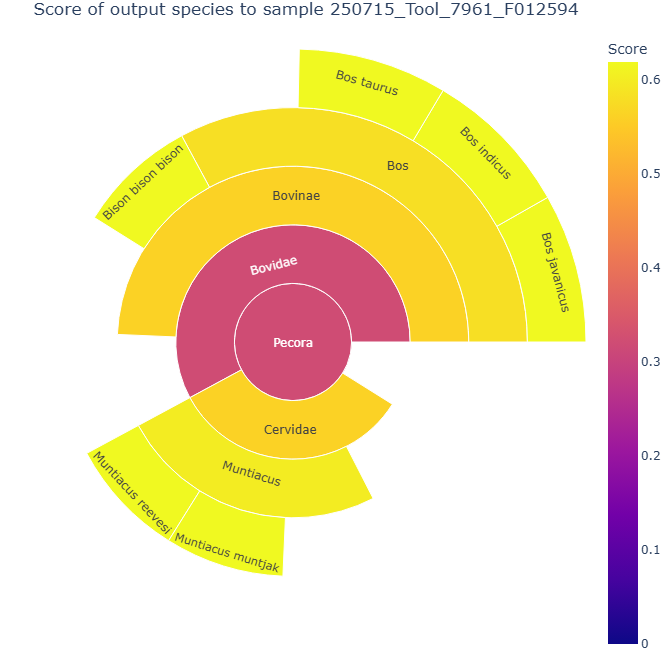

Figure S7: ClassiCOL sunburst outputs below Pecora for tools 8013, 4694, 5258, and 7961.

All ClassiCOL output files (including expanded sunburst plots) are available as supplementary files.

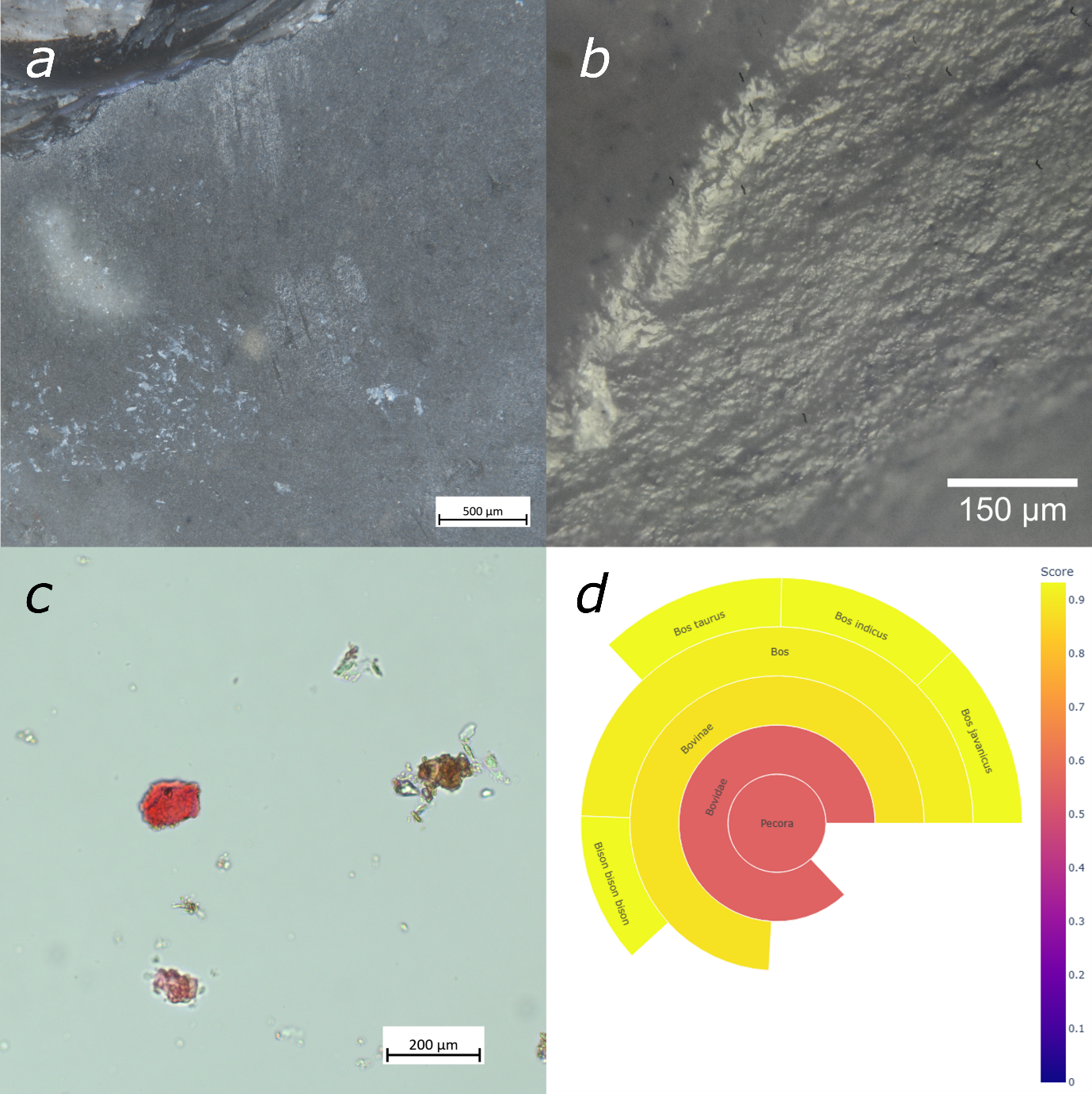

### Extended Discussion

*Figure S8: Multiple analyses of Bazel-Sluis lithic tool 8013 demonstrate consistent evidence for the processing of collagenous animal material: a) white, fatty, amorphous residue smeared onto the surface of the tool, b) well-developed smooth, domed, greasy, bright polish on the very rounded edge of the tool with mixed directionality indicative of contact with bone in a dynamic repetitive motion, c) collagenous tissue stained with Picrosirius red, d) extracted collagen residues are identified as Bovine in a ClassiCOL analysis.*

#### Tool use and processing activities

When considering the likely aims of bone processing activities, the powdering of bone to use as temper seems unlikely as, based on petrographic analysis of the pottery clay, bone-tempered pottery found at Bazel-Sluis was most likely not produced locally (meaning within the Lower-Scheldt basin) (Teetaert, 2020; Teetaert & Crombé, 2022). Processing of bone for marrow extraction, for grease rendering, and for adhesive manufacture remain as possible motivators for bone crushing.

#### On preservation and the wider applicability of lithic proteomics

The state of organic preservation at Bazel-Sluis is very variable according to the location on the site. The best organic preservation is found on the dune slope and borders of the paleochannel; it is from that context that all faunal remains were collected. However, the organic preservation on the dune top, where the lithic industries, faceted tools included, were found needs to be termed bad/poor. Except for calcined bones and charred plant remains (e.g. charcoal, hazelnut shells), there is no organic material preserved. The latter is very common in Western Europe as most Mesolithic/Neolithic sites are situated in dry and acid soils, unfavourable for the preservation of organics. We consider that the fact that at Bazel we could obtain proteomics results on artefacts found in an environment with overall poor organic preservation is very hopeful for future research.

#### Handling of proteomic data and interpretation

In palaeoproteomic datasets, analytical certainty steeply decreases from protein *identification* to species *inference*, to *interpretation* of protein deposition (identification>inference>interpretation). This can be difficult to tackle especially with very sparse datasets and necessitates careful consideration of orthogonal data, such as prior outcomes from artefact analyses and other archaeological finds from the relevant site.

We therefore propose to approach protein identifications on archaeological artefacts in the spirit of jurisdiction: every protein is contamination until proven otherwise. To illustrate this, we consider the proteomic findings from the tools in the framework of an ‘un/certainty matrix’ [figure **S8**]. In this visual representation, every protein starts at the upper right corner (as modern contamination) and moves downward and/or to the left based on increasing evidence.

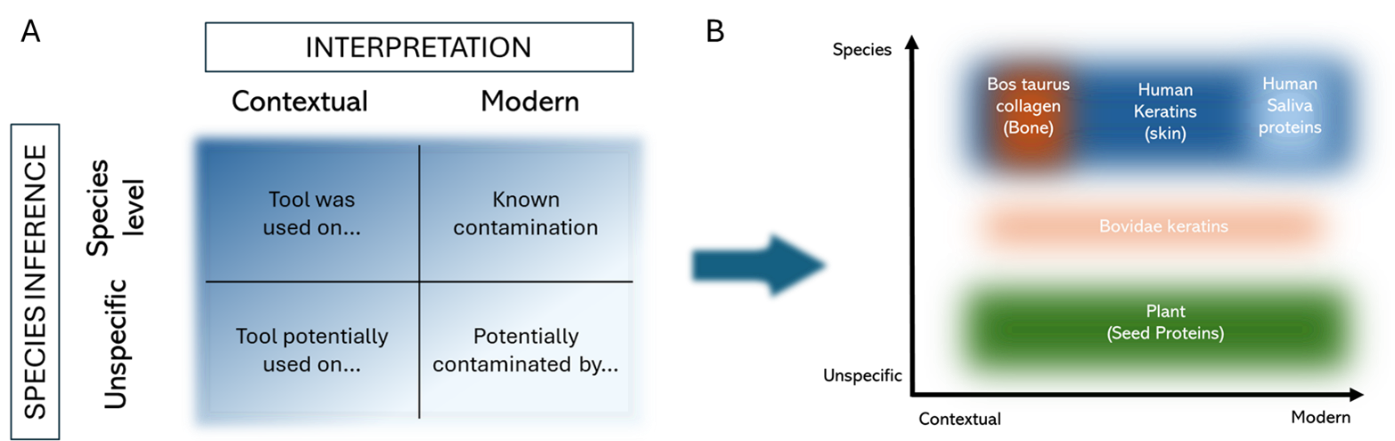

*Figure S9: A proposed framework for classifying archaeological protein identifications at a range of confidence levels. A) an un/certainty matrix for species inference and interpretation based on contextual evidence. B) The most prominent protein identifications from Bazel-Sluis faceted tools placed into an un/certainty matrix. Protein groups unable to be clearly interpretated as contextual or modern span the length of the (temporal) x axis, while proteins whose provenance is considered to be certain occupy small and defined spaces.*

To expand on the placement of proteins within the matrix, we reiterate the following findings. In the case of *Bos taurus* collagen, (i) there are no common sources of bovine collagen contamination in the lab, (ii) collagen is one of the best preserved proteins known to palaeoproteomics, (iii) the use wear analysis indicates that the flint tools were used on bone, (iv) the collagen was identified down to species level by a dedicated species inference tool (ClassiCOL), (v) it was the most prominent protein in the microscopic slides that were stained with picrosirius red and (vi) collagenous-appearing organic trace residues from these tools can be radiocarbon-dated back to the time of residue deposition and implicitly tool use, which matches with other site dating evidence. *Bos taurus* collagens are therefore considered to be a ‘true positive’ resulting from prehistoric tool use, or ‘analytically true, archaeologically true’.

In the case of human keratins, the temporal picture is more muddled, as human skin protein is likely to have been deposited on tools both during use in prehistory, and when the tools were held and examined during and after excavations. Additionally, the preservation of keratins over such long spans of time is not currently as well-evidenced as for protein, with hide, skin and hair not typically identified outside of instances of mummification. Human keratin is also the most common contaminant identified in proteomics datasets, a phenomenon which is even more pronounced in sparse palaeoproteomics data from handled objects. Additionally, keratins were highly abundant in a modern saliva sample, potentially linking (some of) these human keratins to the saliva on the tools. Their identification as human and as keratins, however, is very well-evidenced.

Saliva proteins are identified beyond reasonable doubt through comparison with a modern saliva sample comprised of multiple donations. Their preservation for archaeological timespans is vanishingly unlikely, and paired with extensive anecdotal evidence for preliminary cleaning of tools with saliva during excavations, and of a licked-finger cleaning approach when closely examining artefacts after excavation, the attribution of salivary proteins to a modern origin is also secure.

Animal keratins are ambiguous in both temporal and analytical regards. Firstly, many of the keratin peptides identified can be matched to multiple species, with peptides that do not match to human keratin reference sequences sharing matches to *Ovis* and to *Bos*. Hence these are uncertain taxonomic identifications to the level of the Bovidae family. If these can be attributed to sheep keratins, then it is entirely possible that these relate to modern contact with woollen clothing items; if bovine, these may relate to butchery or to hide processing, as the tool came into contact with skin, hair, and/or horns.

Finally, the plant seed proteins have the lowest level of taxonomic and temporal certainty when considered as a group, as without more comprehensive databases (in themselves requiring better understanding of plant protein preservation), it is not possible to determine the accuracy of some of the species identifications. When taking the identifications at face value, some taxa certainly fit with the archaeological and palynological findings at the site, while others can only represent modern contaminations.
